# Mitochondrial fatty acid synthesis (mtFAS) fuels iron-sulfur cluster production essential for type I immunity

**DOI:** 10.64898/2026.11.24.768974

**Authors:** Samuel E. J. Preston, McLane J. Watson, Lisa M. DeCamp, Michael S. Dahabieh, Joseph Longo, Brandon M. Oswald, Ilaria Panzeri, Susan M. Kitchen-Goosen, Morgan Canfield, Pieter R. Norden, Naomi Graber, Nicole M. Foy, Ryan D. Sheldon, Russell G. Jones, Sara M. Nowinski

**Author notes:** Correspondence: Dr. Sara M. Nowinski,; Dr. Russell G. Jones.

## Abstract

Mitochondria orchestrate the metabolic programs that drive T cell-mediated immunity, yet the specific mitochondrial activities that license T cell effector function remain unclear. Here, we identify mitochondrial fatty acid synthesis (mtFAS) as a critical mitochondrial pathway that supports T cell effector function and limits T cell dysfunction. Upon activation, human and mouse T cells robustly induce mtFAS enzymes in a Myc-dependent manner. Disrupting mtFAS activity in T cells through deletion of the initiating enzyme, malonyl-CoA-acyl carrier protein transacylase (MCAT), sharply reduces effector cytokine production and impairs type I immunity, including CD8^+^ cytotoxic and CD4^+^ Th1 responses, to bacterial and viral pathogens in vivo. Loss of MCAT disrupts protein lipoylation and iron-sulfur cluster (ISC) production in T cells, disrupting the stability and function of electron transport chain (ETC) complexes and impairing oxidative phosphorylation. Deletion of ISC biosynthesis machinery phenocopies MCAT loss, whereas selective disruption of protein lipoylation does not. Moreover, selective restoration of protein lipoylation in MCAT-deficient T cells fails to rescue effector function, establishing ISC biogenesis and its downstream support of ETC activity as the principal mtFAS-dependent mechanism sustaining T cell effector function. MCAT deficiency also promotes the premature acquisition of exhaustion-associated features in CD8^+^ T cells, including increased PD-1 and TIM-3 expression and reduced cytokine competence. Together, these findings establish mtFAS as an essential pathway for initiating and sustaining T cell effector responses, identifying ISC biogenesis and ETC activity as core mtFAS-dependent outputs critical for T cell effector responses and resistance to exhaustion.

## INTRODUCTION

T cells are critical for type I immune defense against pathogens and tumors^1–3^. Upon activation by appropriate antigenic and costimulatory signals, CD8^+^ and CD4^+^ T cells undergo rapid clonal expansion and differentiate into effector T cell (Teff) subsets that produce effector cytokines such as interferon-gamma (IFN-ɣ) and tumor necrosis factor-alpha (TNF-*a*), and in the case of CD8^+^ T cells, gain cytolytic capacity^4,5^. The success of cancer immunotherapies underscores the therapeutic power of robust, sustained T cell effector responses^6^. Yet, persistent antigen stimulation – characteristic in chronic infection and cancer – can drive T cells into a dysfunctional, terminally exhausted state defined by diminished proliferation, attenuated cytokine production, and induction of inhibitory receptors including PD-1, LAG-3, and TIM-3^7–9^. Understanding the mechanisms that direct T cells toward durable effector responses that resist exhaustion is essential for improving immune responses across diverse disease settings.

Metabolic rewiring is fundamental to productive T cell effector function^10,11^. Mitochondria lie at the center of this metabolic transition^12^. In addition to generating ATP through oxidative phosphorylation (OXPHOS), mitochondria are the central hub for a broad array of biosynthetic pathways that are essential for T cell function. These include the tricarboxylic acid (TCA) cycle, which provides both reducing equivalents (i.e., NADH, FADH_2_) and the carbon backbones for amino acid and nucleotide synthesis; electron transport chain (ETC) activity, which produces ATP; and additional anabolic processes that generate acetyl-CoA, aspartate, and pyrimidine nucleotides needed for proliferation^13,14^. Activated T cells require mitochondrial electron transport for full activation and proliferation^15–17^. Notably, exhausted T (Tex) cells exhibit impaired mitochondrial function, reduced OXPHOS, disrupted ETC integrity, and elevated oxidative stress^18–20^. Lowering mitochondrial reactive oxygen species (ROS) can delay terminal exhaustion, suggesting that mitochondrial dysfunction contributes to T cell decline^18,21^. Yet, whether mitochondrial defects initiate loss of effector function or are merely downstream consequences of chronic stimulation remains unresolved. More fundamentally, although mitochondria are required for T cell effector responses^22,23^, the *specific* mitochondrial activities that license T cell effector function and sustain T cell fitness under stress remain unclear.

In addition to housing compartmentalized reactions for central carbon metabolism, mitochondria harbor a dedicated fatty acid synthesis pathway (mtFAS) that is restricted to the mitochondrial matrix and operates independently from cytosolic fatty acid synthesis^24^. This evolutionarily conserved biosynthetic pathway generates specialized acyl chains on an acyl carrier protein (ACP) with two major outputs: (1) octanoyl-ACP conjugates, which fuel lipoic acid synthesis and protein lipoylation, which is required for the function of key enzymes in central carbon metabolism including pyruvate dehydrogenase (PDH) and ɑ-ketoglutarate dehydrogenase (OGDH); and (2) longer chain acyl-ACPs, which stabilize LYRM family assembly factors essential for ETC complex formation and iron–sulfur cluster (ISC) biogenesis^25^. Through these coupled outputs, mtFAS sits at the nexus of oxidative metabolism, coordinating carbon availability with ETC assembly, redox balance, and core mitochondrial metabolic systems. Here, we identify ISC biogenesis and ETC activity as critical mtFAS-dependent outputs required for type I T cell effector function, metabolic fitness, and resistance to exhaustion.

## RESULTS

### T cell activation requires the mtFAS pathway to drive cytokine production and proliferation

Metabolic reprogramming is a fundamental event underlying T cell activation, differentiation, and effector function^26,27^. As mtFAS is a critical regulator of oxidative mitochondrial metabolism (**Fig. 1A**)^25^, we sought to define how mtFAS supports T cell metabolic reprogramming. Data from the ImmPRes database^28^ revealed a robust increase in protein levels of mtFAS pathway machinery, including the acyl carrier protein (ACP), NDUFAB1, and the initiating enzyme malonyl-CoA-acyl carrier protein transacylase (MCAT), as well as downstream targets that require mtFAS products to maintain stability or function (i.e., DLAT, DLST, PDHA1, OGDH, and SDHB), in activated T cells compared to naïve T cells. This upregulation was heavily blunted in cells derived from mice with T cell-specific deletion of *Myc* (*Myc*^fl/fl^;*Cd4*Cre^+^)^29^, a transcription factor and master regulator of T cell metabolic reprogramming^30^, indicating that induction of the mtFAS pathway in both CD8^+^ T cells and CD4^+^ Th1 cells is Myc-dependent (**Fig. 1B**, **Fig. S1A**). RNA sequencing (RNA-seq) of peripheral blood lymphocytes from healthy human donors^31^ revealed a similar increase in expression of 4 key mtFAS components (*MCAT*, *MECR*, *OXSM*, and the gene encoding ACP, *NDUFAB1*) in activated CD8^+^ and CD4^+^ T cells compared to their naïve counterparts (**Fig. 1C**, **Fig. S1B**). Together, these data identify the mtFAS program as an activation-induced metabolic pathway in both human and mouse T cells.

**Figure 1:**
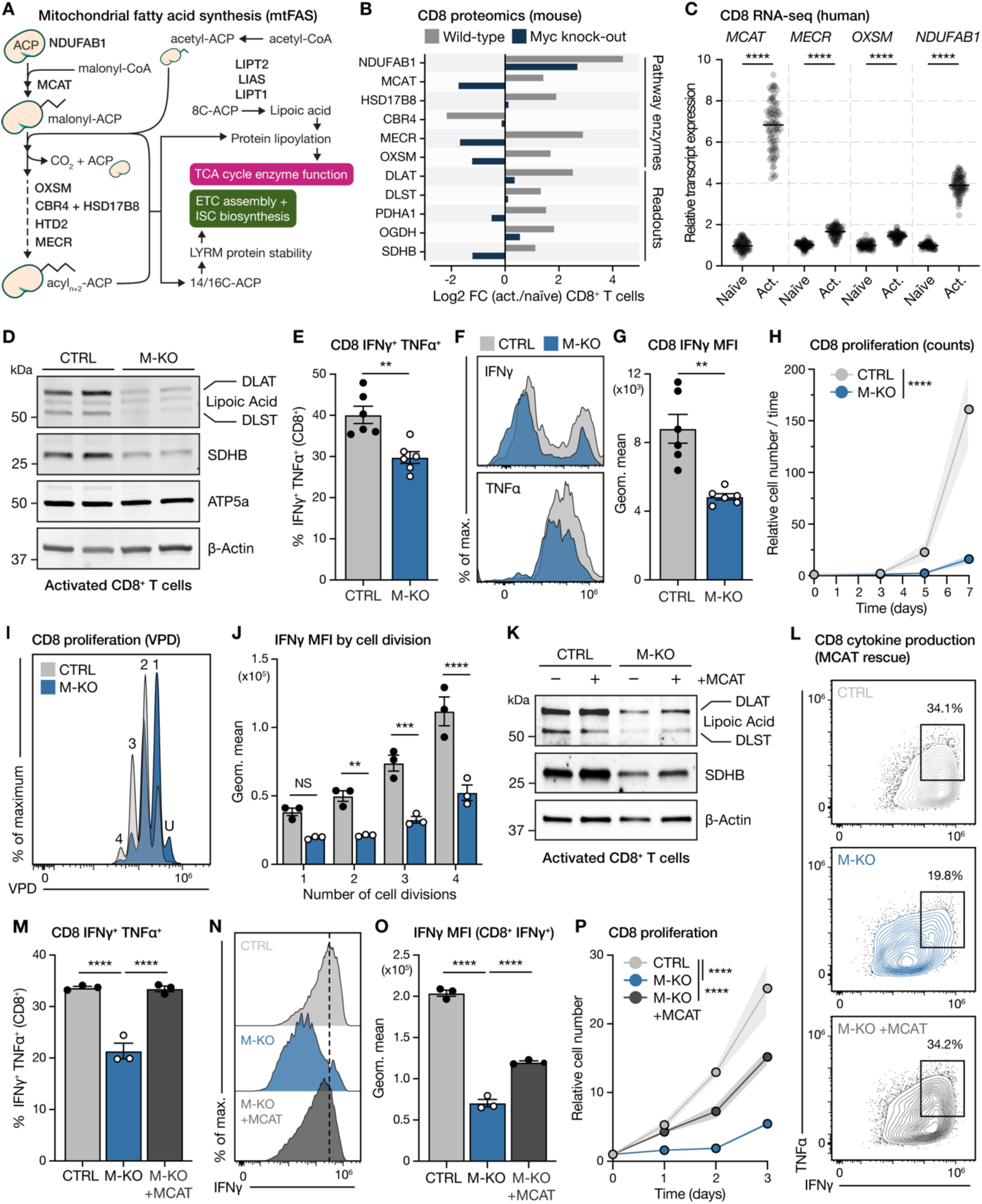
T cell activation requires the mtFAS pathway to drive cytokine production and proliferation. (**A**) Schematic summary of the mitochondrial fatty acid synthesis (mtFAS) pathway. mtFAS enzymes are in bold and how downstream products regulate oxidative mitochondrial metabolism is highlighted. (**B**) Abundance of mtFAS pathway components and downstream mtFAS-dependent proteins in activated CD8^+^ T cells from wild-type and Myc knockout (Myc^fl/fl^;Cd4Cre^+^) mice. Data are from the ImmPRes Myc-regulated T cell proteome dataset and displayed as Log2 fold change in activated cells relative to naïve. (**C**) RNA expression of mtFAS pathway genes in human naïve and activated CD8^+^ T cells isolated from peripheral blood. Data are from the DICE database and shown as transcripts per million. (**D**) Immunoblot of lipoylated DLAT and DLST, SDHB, ATP5a and β-Actin in activated CTRL and M-KO CD8^+^ T cells. (**E-G**) Cytokine production in CTRL and M-KO CD8^+^ T cells activated with plate-bound anti-CD3 and anti-CD28 for 3d, followed by a 4h PMA/ionomycin restimulation prior to analysis. Shown are the percentage of IFN-ɣ^+^ TNF-*a*^+^ CD8^+^ T cells (**E**), representative flow histograms of IFN-ɣ and TNF-*a* expression (**F**) and the mean fluorescence intensity (MFI) of IFN-ɣ levels (**G**). Data represent the mean ± SEM (*n* = 6 mice per group), see also Figure S3B. (**H-I**) Proliferation of CTRL and M-KO CD8^+^ T cells cultured in IL-2 (50 U/mL) following in vitro activation. Shown are the viable T cell counts over 7d of culture (**H**) and representative histograms of violet proliferation dye (VPD) dilution across 72 hours of cell division (**I**). The number of cell divisions are indicated; U denotes undivided cells. Data represent the mean ± SEM (*n* = 3 mice per group), see also Figure S3C. (**J**) MFI of IFN-ɣ levels in CTRL and M-KO CD8^+^ T cells stratified across the number of cell divisions as determined by VPD staining. Data represent the mean ± SEM (*n* = 3 mice per group), see also Figure S3D-G. (**K**) Immunoblot of lipoylated DLAT and DLST, SDHB and β-Actin in activated CTRL and M-KO CD8^+^ T cells with (+) or without (-) expression of murine wild-type MCAT (+MCAT). (**L-O**) Cytokine production in CTRL, M-KO and M-KO +MCAT CD8^+^ T cells. Naïve cells were transduced before activation with plate-bound anti-CD3 and anti-CD28 for 4d, followed by a 4h PMA/ionomycin restimulation prior to analysis. Shown are representative flow plots of IFN-ɣ versus TNF-*a* expression (**L**), the percentage IFN-ɣ^+^ TNF-*a*^+^ cells (**M**), representative histograms of IFN-ɣ expression in CD8^+^ T cells (**N**), and the MFI of IFN-ɣ levels in IFN-ɣ^+^ CD8^+^ T cells (**O**).Data represent the mean ± SD (*n* = 3 technical replicates using cells pooled from 2 mice per genotype). (**P**) Proliferation of CTRL, M-KO and M-KO +MCAT CD8^+^ T cells cultured in IL-2 (50 U/mL). Naïve cells were transduced prior to activation using plate-bound anti-CD3 and anti-CD28 antibodies, then given 3d to recover before assay seeding. The number of viable T cells was then counted over the following 3d. Data represent the mean ± SD (*n* = 3 technical replicates using cells pooled from 2 mice per genotype). For statistics: \*\**p* < 0.01, \*\*\**p* < 0.001, \*\*\*\**p* < 0.0001 by unpaired *t* test with Welch’s correction (**E and G**), one-way ANOVA with Tukey’s multiple comparisons test (**C, J, M and O**), or two-way ANOVA with Tukey’s multiple comparisons test (**H, J and P**).

Given our observations comparing naïve and activated T cells, we next examined the contribution of mtFAS activity to the metabolic changes that underpin T cell effector function. To investigate this, we generated mice with specific loss of mtFAS activity in mature T cells by crossing *CD4*^Cre^ transgenic mice with mice harboring floxed alleles of the *Mcat* gene (*Mcat*^fl/fl^)^32^. Biochemical analysis of activated T cells from control (denoted CTRL, *Mcat*^fl/fl^;*Cd4*Cre^-^) and T cell MCAT-deficient (denoted M-KO, *Mcat*^fl/fl^; C*d4*Cre^+^) mice revealed a significant reduction in both MCAT protein (**Fig. S1C**) and *Mcat* mRNA (**Fig. S1D**) in M-KO CD8^+^ T cells and CD4^+^ Th1 cells. Downstream functional targets of mtFAS, including lipoylation of PDH (DLAT) and OGDH (DLST) and stabilization of succinate dehydrogenase B (SDHB), a surrogate for the ETC complex assembly function of mtFAS products^33^, were reduced in M-KO CD8^+^ and CD4^+^ Th1 cells compared to controls, indicating reduced mtFAS activity in mature T cells using this targeting strategy (**Fig. 1D**, **Fig. S1E**). MCAT loss had minimal effect on overall T cell homeostasis in mature mice (8-10 weeks of age). Overall splenocyte and T cell numbers were equal between control and M-KO animals (**Fig. S1F-G**), and we observed no significant difference in the proportion of CD8^+^ versus CD4^+^ T cells in the spleens of CTRL versus M-KO mice (**Fig. S1H-K**). Moreover, there was no significant difference in the proportion of CD44^+^ or CD62L^+^ T cell subsets at baseline (**Fig. S1L-S**), with the exception of the CD44^+^CD62L^-^ CD8^+^ and CD4^+^ T effector memory (Tem) cell populations, which were significantly lower in M-KO mice (**Fig. S1N,R**). Thus, MCAT loss does not appear to alter the development or homeostasis of naïve T cells.

We next tested the role of mtFAS in T cell effector responses in vitro. CD8^+^ T cells isolated from CTRL and M-KO mice activated normally in response to anti-CD3 and anti-CD28 antibody stimulation as evidenced by similar upregulation of CD44, CD25, CD69, T-Bet, and PD-1 expression up to 72 hours post-activation (**Fig. S2A-E**). However, MCAT deletion robustly hindered CD8^+^ T cell effector function, as M-KO CD8^+^ T cells displayed reduced expression of Granzyme B post-activation (**Fig. S3A**) and produced significantly less IFN-ɣ and TNF-*a* compared to CTRL CD8^+^ T cells (**Fig. 1E-G**, **Fig. S3B**). Proliferation was also markedly impacted by MCAT deletion, with M-KO CD8^+^ T cells displaying reduced expansion (**Fig. 1H**) and undergoing fewer cell divisions over time (**Fig. 1I**, **Fig. S3C**) following anti-CD3 and anti-CD28 stimulation. To test whether the observed reduction in IFN-ɣ production could be explained by this proliferative defect, we examined IFN-ɣ production across cell divisions and found that IFN-ɣ levels were lower in M-KO T cells at each cell division (**Fig. 1J**, **Fig. S3D-G**). In contrast, T-Bet expression was uniform between CTRL and M-KO CD8^+^ T cells across all cell divisions (**Fig. S3H-I**). These data suggest that mtFAS loss does not broadly impair T cell differentiation, but rather impacts cytokine production. Re-introducing wild type mouse MCAT into M-KO CD8^+^ T cells rescued protein lipoylation and SDHB stability (**Fig. 1K**), cytokine production (**Fig. L-O**), and cell proliferation (**Fig. 1P**).

Analysis of MCAT-deficient CD4^+^ T cells revealed similar defects in cytokine production as M-KO CD8^+^ T cells. CD4^+^ T cells were activated in vitro and polarized to a Th1 subtype by the addition of IL-12 to their media (herein referred to as CD4^+^ Th1 cells). Similar to CD8^+^ T cells, cytokine production was significantly reduced in M-KO CD4^+^ Th1 cells compared to control cells (**Fig. S4A-C**), despite similar levels of activation (**Fig. S4D**). In contrast, CD4^+^ Th1 cell proliferation was largely unaffected by MCAT loss, although CD4^+^ Th1 cell growth rates overall were slower compared to CD8^+^ T cells (**Fig. S4E-G**). The effect of MCAT loss on CD4^+^ T cell effector function appeared to be limited to the Th1 lineage, as only Th1 cell polarization was impaired by *Mcat* deletion (**Fig. S4H**). Analysis of ImmPRes proteomics data indicated that Th1 cells have higher protein levels of mtFAS machinery and pathway readouts compared to Th2, Th17 and Treg cells (**Fig. S4I**), supporting the specific effect of MCAT loss on Th1 cell function. Combined, our data reveal that MCAT loss blunts cytokine production in CD8^+^ and CD4^+^ Th1 T cells, key mediators of type I immunity.

### mtFAS is essential for type I immune responses in vivo

Given the reliance of CD8^+^ and CD4^+^ Th1 T cell effector function on mtFAS activity, we next assessed the impact of mtFAS deficiency on type I immune responses to bacterial and viral infection in vivo. We first assessed CD8^+^ and CD4^+^ Th1 T cells in CTRL versus M-KO mice in response to infection with an attenuated strain of *Listeria monocytogenes* expressing ovalbumin (LmOVA) (**Fig. 2A**). At 7 days post-infection (dpi), when the antibacterial T cell response is at its peak, we observed significantly fewer CD8^+^ and CD4^+^ Th1 T cells in the spleens of M-KO mice, and these T cells were less activated (CD44^hi^) compared to their CTRL counterparts (**Fig. 2B-E**). To assess antigen-specific cytokine production, we cultured splenocytes isolated from LmOVA-infected mice with either OVA_257-264_ (SIINFEKL) or LLO_190-201_ peptides to stimulate CD8^+^ and CD4^+^ Th1 cells, respectively. While CTRL mice displayed normal IFN-ɣ and TNF-*a* production by CD8^+^ and CD4^+^ Th1 T cells, cytokine production by M-KO T cells was completely ablated (**Fig. 2F-I**).

**Figure 2:**
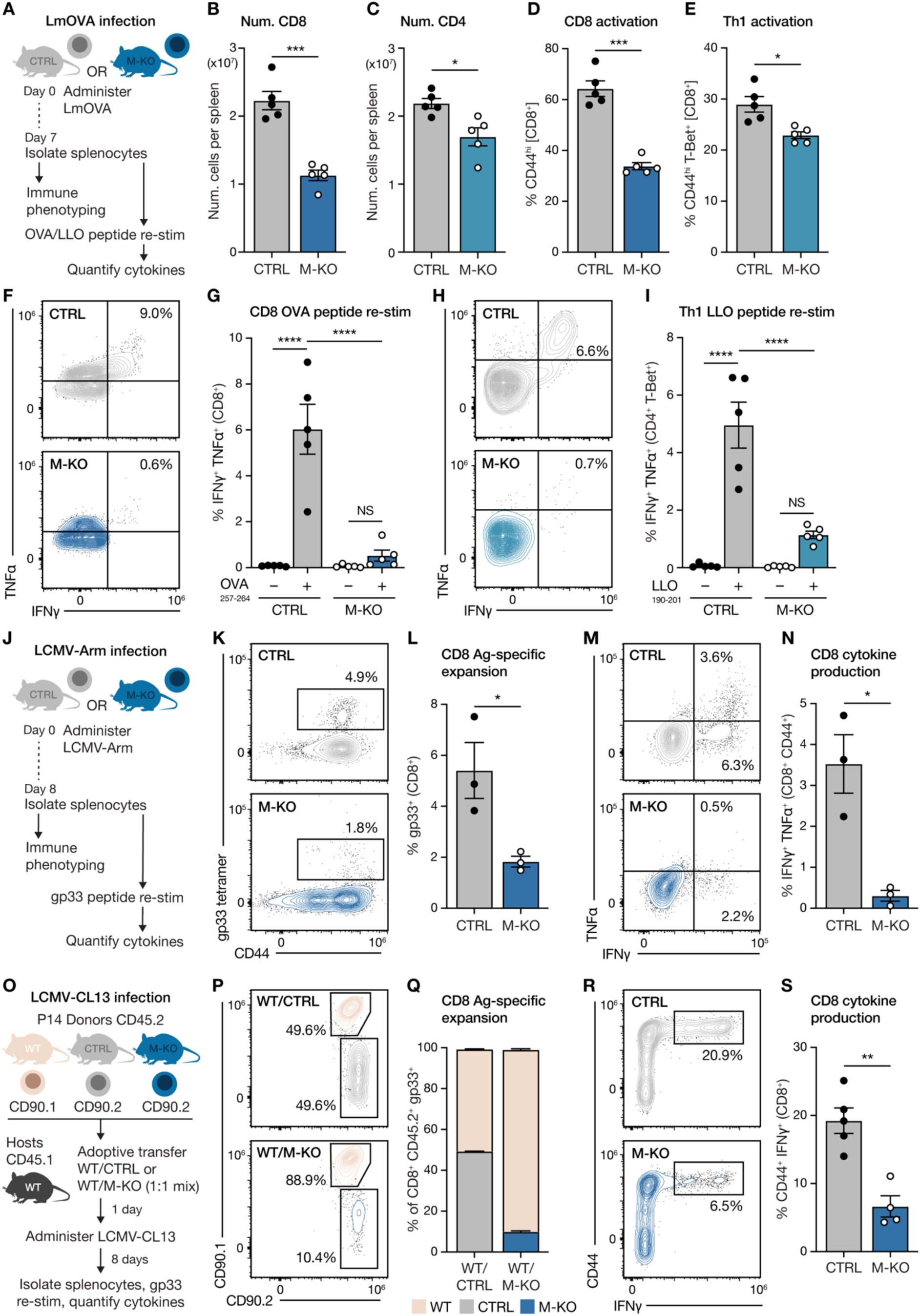
mtFAS is essential for type I immune responses in vivo. (**A**) Experimental setup for *Listeria monocytogenes* (LmOVA) infection of CTRL or M-KO mice. (**B-E**) Total number of CD8^+^ (**B**) and CD4^+^ (**C**) T cells and the percentage of activated (CD44^+^) CD8^+^ T (**D**) and CD4^+^ Th1 (**E**) cells the in the spleens of CTRL and M-KO mice at 7 days post-LmOVA infection (dpi). Data represent the mean ± SEM (*n* = 5 mice per group). (**F-I**) Antigen-specific cytokine responses by CD8^+^ (**F-G**) and CD4^+^ (**H-I**) T cells from CTRL and M-KO mice following LmOVA infection (7 dpi). Shown are representative flow cytometry plots for IFN-ɣ versus TNF-*a* expression (**F and H**) and quantification of the percentage of IFN-ɣ^+^TNF-*a*^+^ T cells (**G and I**) from infected CTRL and M-KO mice at 7 dpi. Cytokine production in CD8^+^ and CD4^+^ T cells was measured following *ex vivo* culture for 4h with or without OVA (**G**) or LLO (**I**) peptide. Data represent the mean ± SEM (*n* = 5 mice per group). (**J**) Experimental setup for infection of CTRL or M-KO mice with LCMV Armstrong strain (LCMV-Arm). (**K-L**) Expansion of antigen-specific CD8^+^ T cells following LCMV infection (8 dpi). Representative flow cytometry plots for gp33 versus CD44 levels (**K**) and quantification of the percentage of gp33^+^ CD8^+^ T cells in the spleens of CTRL versus M-KO mice following LCMV-Arm infection (8 dpi). Data represent the mean ± SEM (*n* = 3 mice per group). (**M-N**) IFN-ɣ versus TNF-*a* production by antigen-specific CD8^+^ T cells responding to LCMV-Arm. Shown are representative flow cytometry plots (**M**) and quantification of the percentage of IFN-ɣ^+^TNF-*a*^+^ CD44^+^ CD8^+^ T cells (**N**) from LCMV-Arm infected CTRL and M-KO mice at 8 dpi. Cytokine production in CD8^+^ T cells was measured following *ex vivo* culture for 4h with gp33 peptide. Data represent the mean ± SEM (*n* = 3 mice per group). (**O**) Experimental setup for the co-adoptive transfer of WT CD90.1^+^ P14 cells with CTRL or M-KO CD90.2^+^ CD8^+^ P14 cells into congenic hosts, followed by infection with LCMV Clone 13 strain (LCMV-CL13). (**P-Q**) Expansion of CTRL versus M-KO P14 T cells in vivo following LCMV-CL13 infection. Shown are representative flow cytometry plots for CD90.1^+^ versus CD90.2^+^ adoptively transferred P14 T cells of the indicted genotypes 8 dpi with LCMV-CL13 (**P**) and quantification of the proportion of WT CD90.1^+^ versus CTRL or M-KO CD90.2^+^gp33^+^ CD8^+^ P14 cells in the spleens of host mice at 8 dpi (**Q**). Data represent the mean ± SEM (*n* = 5 for WT/CTRL, 4 for WT/M-KO). (**R-S**) Representative flow cytometry plots for IFN-ɣ versus CD44 expression by P14 T cells (**R**) and quantification of the proportion of CD44^+^ IFN-ɣ^+^ CTRL and M-KO CD90.2^+^ CD8^+^ P14 cells (**S**) from the spleens of host mice at 8 dpi with LCMV-CL13. Cytokine production by P14 T cells of each genotype was measured following *ex vivo* culture for 4h with gp33 peptide. Data represent the mean ± SEM (*n* = 5 WT/CTRL, 4 WT/M-KO). For statistics: \**p* > 0.05, \*\**p* < 0.01, \*\*\**p* < 0.001, \*\*\*\**p* < 0.0001 by unpaired *t* test with Welch’s correction (**B-E, L, N and S**), or one-way ANOVA with Tukey’s multiple comparisons test (**G and I**), NS denotes not significant.

As the T cell response to LmOVA infection is largely CD8^+^ T cell-mediated, we used *Citrobacter rodentium* to model a T cell response to bacterial infection in the large intestine that is primarily CD4^+^ T cell-driven (i.e., Th1 and Th17 cell)^34^ (**Fig. S5A**). We observed no difference in whole-body response to *Citrobacter* infection between CTRL and M-KO mice, as measured by body weight changes, large intestine weight, and large intestine length (**Fig. S5B-D**). Despite this, quantifying *C. rodentium* bacterial content in the feces of infected mice over time revealed that M-KO mice had an impaired ability to clear their infection (**Fig. S5E-G**). Analysis of T cell populations in the lamina propria of the large intestine at 7 dpi revealed a modest reduction in the proportion of CD4^+^ T cells (**Fig. S5H**) but not their activation (**Fig. S5I**). These CD4^+^ T cells produced significantly less IFN-ɣ and TNF-*a* than controls (**Fig. S5J-L**), with their impaired function likely contributing to reduced infection control.

To assess the impact of T cell mtFAS on antiviral immunity, we quantified CD8^+^ T cell responses to lymphocytic choriomeningitis virus (LCMV) infection. We first assessed CD8^+^ T cell responses to acute viral infection in CTRL and M-KO mice following systemic infection with LCMV-Armstrong strain (LCMV-Arm) (**Fig. 2J**). At 8 dpi, we observed a dramatic reduction in the expansion of LCMV-specific gp33^+^ CD8^+^ T cells in M-KO mice compared to controls (**Fig. 2K-L**). Similar to our observations in LmOVA infection (**Fig. 2F-G**), cytokine production by CD8^+^ T cells was ablated by MCAT deletion, as quantified by the percentage of IFN-ɣ^+^ TNF-*a*^+^ CD8^+^ T cells in the spleen of infected M-KO animals at 8 dpi (**Fig. 2M-N**). To establish whether the effect of MCAT loss on CD8^+^ T cell expansion and effector function is T cell-intrinsic, we co-transferred equal numbers of P14 CD90.1^+^ wild-type (WT) CD8^+^ T cells with either CTRL or M-KO P14 CD90.2^+^ CD8^+^ T cells into congenic hosts, followed by infection with LCMV-Clone13 strain (LCMV-CL13) (**Fig. 2O**). At 8 dpi, we observed an equal expansion of LCMV-specific gp33^+^ P14 CTRL and WT CD8^+^ T cells; in contrast, the expansion of MCAT-deficient P14 CD8^+^ T cells was markedly reduced compared to WT P14 T cells (**Fig. 2P-Q**), even in the presence of T cells capable of responding to LCMV. Consistent with data seen in M-KO animals (**Fig. 2M-N**), these P14 M-KO CD8^+^ T cells produced significantly less IFN-ɣ on a per-cell basis than CTRL P14 cells (**Fig. 2R-S**). Collectively, our findings confirm that mtFAS is required to fuel antigen-specific CD8^+^ and CD4^+^ Th1 T cell expansion and cytokine production in response to bacterial and viral pathogen infection in vivo.

### MCAT loss impairs T cell oxygen consumption and bioenergetic capacity

Given the impaired effector function of mtFAS-deficient T cells, we next investigated the molecular consequences downstream of *Mcat* deletion in T cells. As the mtFAS pathway programs oxidative mitochondrial metabolism by supporting TCA and ETC function^33,35^, we assessed the bioenergetics of CTRL and M-KO CD8^+^ and CD4^+^ Th1 cells using a Seahorse XF96 Extracellular Flux Analyzer. Both CD8^+^ T cells and CD4^+^ Th1 cells with MCAT deletion displayed reduced basal and maximal oxygen consumption rates (OCR), indicating reduced respiratory capacity (**Fig. 3A**, **Fig. S6A-C**). Interestingly, the reduction in OCR in M-KO T cells was not compensated by an increase in glycolysis as determined by a reduced extracellular acidification rate (ECAR) (**Fig. 3B**, **Fig. S6D**), suggesting an overall blunting of bioenergetic output in mtFAS-deficient T cells.

**Figure 3:**
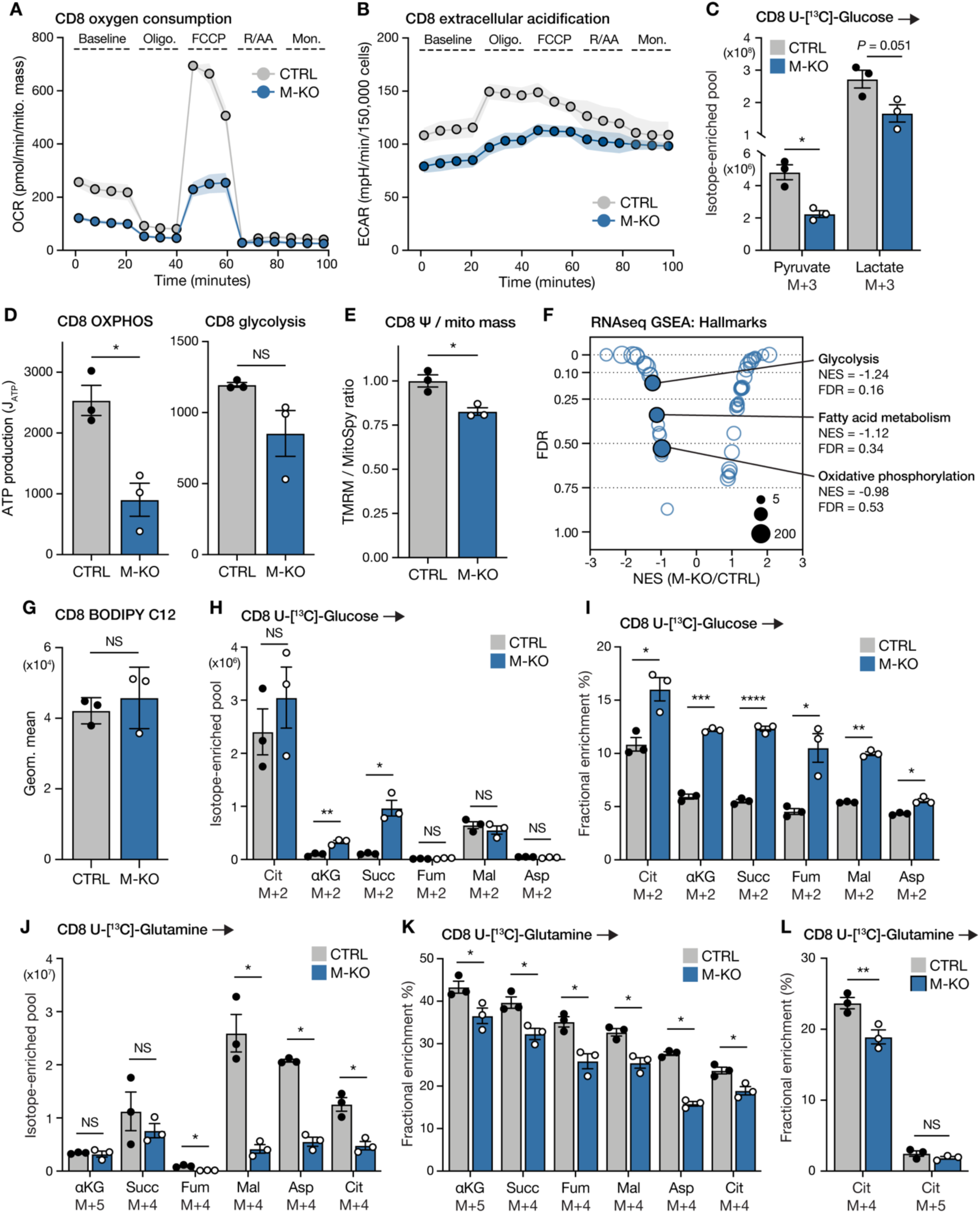
MCAT loss impairs T cell oxygen consumption and bioenergetic capacity. (**A and B**) Oxygen consumption rate (OCR, **A**) and extracellular acidification rate (ECAR, **B**) over time of activated CTRL and M-KO CD8^+^ T cells, as quantified using the Seahorse mitochondrial stress test. OCR data are normalized to cellular mitochondrial content (**A**, see also Figure S6A for per-cell normalization). Data show one representative sample per genotype from 3 independent biological replicate, error bars denote SD. (**C**) Contribution of [U-^13^C]-glucose to pyruvate and lactate in activated CTRL and M-KO CD8^+^ T cells. Data represent the total abundance of the indicated metabolite mass isotopologues derived from labeled glucose; error bars denote SEM (*n* = 3 mice per group). (**D**) Calculated maximal ATP production rates derived from oxidative phosphorylation (OXPHOS) and glycolysis in CTRL and M-KO CD8^+^ T cells. Data represent the mean ± SEM (*n* = 3 mice per group). (**E**) Quantification of mitochondrial membrane potential (Ψ) normalized to cellular mitochondrial content in CTRL and M-KO CD8^+^ T cells. Data represent the mean ± SEM (*n* = 3 mice per group), see also Figure S6G for individual MitoSpy and TMRM stainings. (**F**) Gene set enrichment analysis (GSEA) showing GO terms that are enriched in M-KO versus CTRL CD8+ T cells based on RNA transcript levels (*n* = 6 mice per group). Bubbles display the normalized enrichment score (NES) and false discovery rate (FDR) for all analyzed signatures in the MSigDB Hallmarks database; bubble size equates to gene list size. Select negatively-enriched are highlighted alongside their NES and FDR values. See also Figure S6I-J. (**G**) Quantification of BODIPY C12 (fluorescent fatty acid analog) uptake in activated CTRL and M-KO CD8^+^ T cells following a 30m incubation. Data represent the mean ± SEM (*n* = 3 per group). (**H-L**) Contribution of [U-^13^C]-glucose (**H and I**) and -glutamine (**J-L**) to TCA cycle intermediates and aspartate in activated CTRL and M-KO CD8^+^ T cells following a 4h trace. Data shown represent the total abundance of the indicated metabolite mass isotopologues derived from the labeled tracer (**H and J**), or fractional enrichment of the labeled tracer in the indicated mass isotopologues (**I, K and L**). Error bars denote SEM (*n* = 3 mice per group per tracer). For statistics: \**p* > 0.05, \*\**p* < 0.01, \*\*\**p* < 0.001, \*\*\*\**p* < 0.0001 by unpaired *t* test with Welch’s correction (**C-E, G-K**), or two-way ANOVA with Tukey’s multiple comparisons test (**L**), NS denotes not significant.

Consistent with this, ^13^C-glucose tracing revealed decreased labeling of pyruvate and lactate from glucose in M-KO CD8^+^ T cells compared with controls (**Fig. 3C**). Calculated ATP production rates from both OXPHOS and glycolysis were significantly lower in both M-KO CD8^+^ T cells and CD4^+^ Th1 cells compared to controls, with ATP production from OXPHOS displaying the strongest dependence on MCAT (**Fig. 3D**, **Fig. S6E**). This correlated with an overall reduction in mitochondrial membrane potential (Ψ) in M-KO T cells, as measured by the ratio of TMRM to MitoSpy fluorescence to account for differences in cellular mitochondrial content (**Fig. 3E**, **Fig. S6F**). Interestingly, we observed that mitochondrial mass was increased in T cells lacking MCAT, without an equivalent increase in membrane potential (**Fig. S6G-H**). To assess whether impaired glucose-dependent metabolism and OXPHOS might be compensated by increased fatty acid oxidation in M-KO cells, we performed bulk RNA-seq on activated CTRL and M-KO CD8^+^ T cells (**Fig. S6I-J**). Gene set enrichment analysis (GSEA) identified fatty acid metabolism as negatively enriched in M-KO cells compared to controls, in addition to glycolysis and OXPHOS signatures, consistent with our bioenergetic profiling (**Fig. 3F**). We further performed BODIPY C12 labeling assays to functionally assess fatty acid uptake and found no difference in BODIPY fluorescence between CTRL and M-KO CD8^+^ T cells (**Fig. 3G**). Taken together, these data suggest that *Mcat*-deleted T cells actively upregulate mitochondrial biogenesis, but without restoring mtFAS activity these new mitochondria remain dysfunctional.

As discussed above, in addition to supporting ETC function, the mtFAS pathway maintains the activity of several key TCA cycle-related enzymes, including PDH, OGDH, and SDH. We hypothesized that decreased TCA cycle activity may account for the reduction in oxidative ATP production in MCAT-deficient T cells. Using metabolomics to track the incorporation of uniformly labeled ^13^C-glucose or ^13^C-glutamine into TCA cycle intermediates and aspartate, we found that MCAT loss has differing effects on TCA cycle metabolism in CD8^+^ T cells versus CD4^+^ Th1 T cells. M-KO CD8^+^ T cells displayed ^13^C-labeling patterns consistent with decreased SDH activity, marked by a significant increase in the abundance of ^13^C-glucose-derived alpha-ketoglutarate and succinate (m+2), while labeling into downstream metabolites was unchanged relative to CTRL CD8^+^ T cells (**Fig. 3H-I**). Similarly, when tracing with ^13^C-glutamine, M-KO CD8^+^ T cells displayed significantly reduced labeling into metabolites downstream of SDH: fumarate, malate, aspartate, and citrate (m+4) (**Fig. 3J-K**). In contrast, the contribution of ^13^C-glucose or ^13^C-glutamine to TCA cycle metabolites in M-KO CD4^+^ Th1 cells was comparable to Th1 cells with an intact mtFAS pathway (**Fig. S7A-D**). One potential impact of ETC dysfunction on the TCA cycle is a metabolic switch to reductive carboxylation of glutamine to sustain citrate and cytosolic acetyl-CoA production for biosynthesis^36,37^. Despite reduced mitochondrial membrane potential and reduced oxidative capacity, MCAT loss in either CD8^+^ T cells or CD4^+^ Th1 cells did not induce reductive carboxylation, as indicated by low ^13^C-glutamine labeling into citrate m+5 (**Fig. 3L**, **Fig. S7E**). Overall, these results indicate that mtFAS regulates the overall bioenergetic capacity of T cells.

### Disrupted complex I formation and iron-sulfur cluster biosynthesis are central features of mtFAS deficiency in T cells

The fact that MCAT loss reduces effector function and OXPHOS in both CD8^+^ and CD4^+^ T cells yet exerts divergent effects on TCA cycle metabolism implies that mtFAS controls a common regulatory pathway in T cells. To delineate the molecular mechanisms linking mtFAS to T cell effector function, we performed global proteomic profiling of activated CTRL and M-KO CD8^+^ T cells. GSEA of the resulting data identified distinct signatures associated with OXPHOS, ETC complex function, and iron-sulfur cluster (ISC) biogenesis that were lost in M-KO CD8^+^ T cells compared to control T cells (**Fig. 4A**). We performed leading edge analysis on all significantly downregulated GO terms and pathways to determine the degree of overlap between these signatures and identify the strongest contributors to their enrichment score, thereby defining the core set of proteins altered by mtFAS deficiency (**Fig. 4B-C**). We observed a group of 34 ETC proteins, predominantly from ETC complex I (CI), and a group of 27 ISC-related proteins, including 5 ISC-containing CI subunits, that lead their respective signatures (**Fig. 4D**). These findings were consistent with those from an equivalent proteomic profiling experiment performed on CD4^+^ Th1 cells, which found that CI formation and ISC biosynthesis (detected within a complex III assembly gene set) were the only pathways significantly reduced by MCAT loss in CD4^+^ T cells (**Fig. S8A-C**). While glycolysis, cholesterol biosynthesis, and nucleotide metabolism signatures were also downregulated in M-KO CD8^+^ T cells (**Fig. 4A-C**), these changes were not observed in Th1 cells (**Fig. S8A-B**). Consistent with a complex I-driven phenotype, we saw an elevation of the lactate:pyruvate ratio in M-KO CD8^+^ T cells compared to control T cells, which is an indicator of a decreased NAD^+^:NADH ratio (**Fig. S8D**). Therefore, for subsequent analyses, we focused on CI assembly/stability and ISC biosynthesis as pathways commonly dysregulated by MCAT loss in CD8^+^ T and CD4^+^ Th1 cells that may link mtFAS activity to T cell effector function.

**Figure 4:**
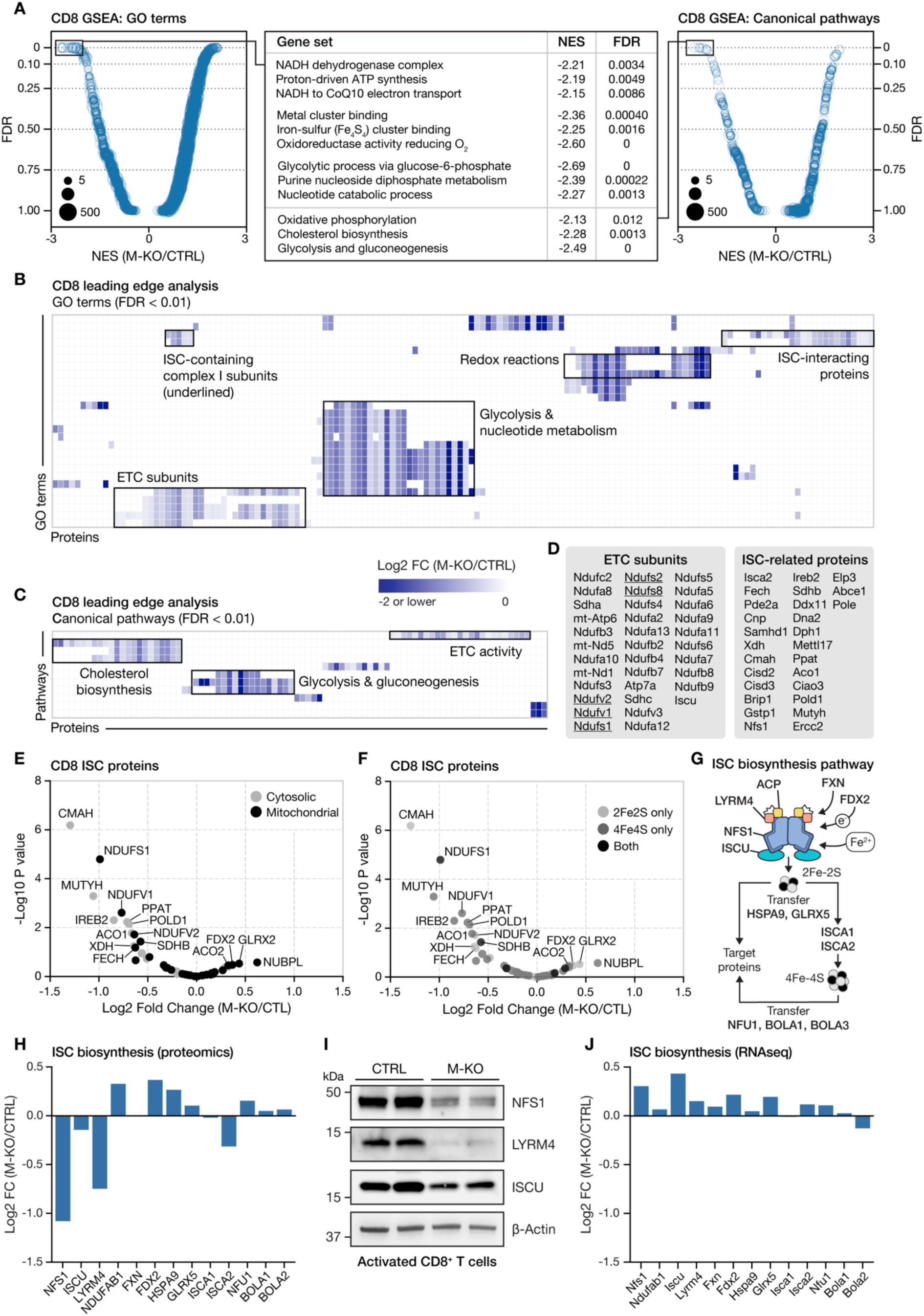
Disrupted complex I formation and iron-sulfur cluster biosynthesis are central features of MCAT-knockout T cells. (**A**) Gene set enrichment analysis (GSEA, MSigDB C2 and C5) showing GO terms and canonical pathways that are enriched in M-KO versus CTRL CD8^+^ T cells based on total protein abundance. Bubble plots display the normalized enrichment score (NES) and false discovery rate (FDR) for all analyzed signatures; bubble size equates to gene list size. A selection of negatively-enriched signatures is listed (middle table) alongside their NES and FDR values. (**B and C**) Heatmap of protein levels (log2 fold change, M-KO/CTRL) for leading edge proteins in all signatures with significant negative enrichment in M-KO CD8^+^ T cells compared to CTRL CD8^+^ T cells. (**D**) List of electron transport chain (ETC) subunits and iron-sulfur cluster (ISC) proteins identified by leading edge analysis in **B**. ISC-containing complex I subunits are underlined. (**E and F**) Volcano plots displaying the protein levels of known ISC proteins in CTRL and M-KO CD8^+^ T cells. Protein abundance is shown relative to CTRL (log2 fold change, M-KO/CTRL); dot color denotes cytosolic versus mitochondrial localization (**E**) or type of ISC involved (**F**). (**G**) Schematic of the ISC biosynthesis pathway. (**H**) Waterfall plot displaying the protein level of ISC biosynthesis machinery in CTRL and M-KO CD8^+^ T cells. Data are expressed as relative to CTRL (combined from *n* = 4 CTRL, 3 M-KO). (**I**) Immunoblot of NFS1, LYRM4, ISCU and β-Actin protein levels in activated CTRL and M-KO CD8^+^ T cells. (**J**) Waterfall plot displaying the RNA transcript levels of ISC biosynthesis machinery in CTRL and M-KO CD8^+^ T cells. Data are expressed as relative to CTRL (combined from *n* = 6 per group

To test whether loss of ETC complex I activity was the major driver of impaired T cell effector function in M-KO mice, we crossed our mice with Rosa26-LSL-NDI1 transgenic knock-in mice^38^ to create MCAT-deficient T cells with ectopic expression of NDI1 (M-KO +NDI1 mice). *Ndi1* is a *Saccharomyces cerevisiae* gene that encodes an alternative NADH dehydrogenase which transfers electrons from NADH to coenzyme Q, functioning similarly to mammalian complex I but without pumping protons. Expression of NDI1 has been shown to rescue specific complex I loss in a variety of models^38–41^. Indeed, expression of NDI1 increased the basal and maximal OCR of M-KO CD8^+^ T cells (**Fig. S8E**). Despite this, NDI1 failed to rescue the proliferation or cytokine production of MCAT-deficient CD8^+^ T cells (**Fig. S8F-J**).

Further investigation of our proteomics data revealed which ISC proteins were most affected by the loss of mtFAS: the three CI subunits NDUFS1, NDUFV1, and NDUFV2 were consistently reduced in MCAT-deficient CD8^+^ and CD4^+^ Th1 T cells (**Fig. 4E-F**, **Fig. S8K-L**). Conversely, levels of the ISC assembly factor NUBPL, which helps to transfer ISCs into CI subunits during assembly^42^, was increased, potentially indicating a compensatory mechanism to increase CI function in T cells when *Mcat* is deleted (**Fig. 4E-F**, **Fig. S8K-L**). Levels of IREB2, which regulates cellular iron homeostasis via transcription of iron-responsive genes, were also lower in M-KO CD8^+^ T cells, further suggesting that intracellular iron levels may be imbalanced (**Fig. 4E-F**). The ISC proteins downregulated by MCAT loss were not stratified by cytosolic versus mitochondrial subcellular localization (**Fig. 4E**, **Fig. 8K**) nor by the type of ISC involved (**Fig. 4F**, **Fig. S8L**). We therefore hypothesized that these combined effects were a downstream consequence of disrupted ISC biogenesis.

The core ISC biogenesis complex comprises NFS1, ISCU, LYRM4, and the mitochondrial acyl carrier protein (ACP, *NDUFAB1*). Stability and function of the complex requires the physical interaction of LYRM4 with mtFAS products attached to ACP^33,43,44^. NFS1, LYRM4, and ISCU were some of the strongest down-regulated proteins identified by our leading edge analyses (**Fig. 4B-D**, **Fig. S8A-C**). In addition to these core proteins, the ISC complex also contains the regulatory protein FXN and electron donor FDX2. This multimer creates 2Fe-2S clusters which are then transferred by HSPA9, GLRX5, and other accessory units to their destination proteins, or to ISCA1 and ISCA2 for the formation of 4Fe-4S clusters. Mature clusters are then inserted into their target proteins by several factors, including NFU1, BOLA1 and BOLA2 (pathway summarized in **Fig. 4G**). NFS1, LYRM4 and ISCU were the only components of this machinery that were downregulated in both M-KO CD8^+^ T cells and CD4^+^ Th1 cells, as measured by proteomics (**Fig. 4H**, **Fig. S8M**) and confirmed by immunoblot (**Fig. 4I**, **Fig. S8N**). Notably, transcript levels for *Nfs1*, *Lyrm4*, and *Iscu* were unchanged in M-KO compared to control CD8^+^ T cells via RNA-seq analysis (**Fig. 4J**). These data support the notion that ISC protein levels in M-KO T cells result from the post-translational stabilization of the ISC biogenesis complex, as has been shown in other models^33,43,44^. Thus, our combined proteomics and bioenergetics data (**Fig. 3A-E**) suggest that MCAT loss disrupts formation of the mitochondrial core ISC complex in T cells, thereby reducing ISC production and compromising downstream CI activity.

### mtFAS supports effector T cell function via regulation of ISC synthesis

We next asked how mtFAS-dependent control of the NFS1-LYRM4-ISCU core ISC production complex impacts T cell proliferation and effector function. We first assessed whether disruption of mtFAS-mediated ISC biogenesis changes T cell iron dynamics. When culturing CTRL and M-KO T cells, we observed that pellets of *Mcat*-deficient CD8^+^ T cells and CD4^+^ Th1 cells were darker than controls (**Fig. 5A**, **Fig. S9A**), suggesting increased iron content. To test this, we used fluorescent probes to quantify the abundance of free ferrous (Fe^2+^) iron in live cells. At a whole-cell level, we found that MCAT loss disrupted iron homeostasis but in opposing directions between CD8^+^ and Th1 cells; M-KO CD8^+^ T cells had significantly more free iron per cell than control cells (**Fig. 5B-C**), while M-KO Th1 had less (**Fig. S9B-C**). However, free iron levels *within mitochondria* were significantly elevated in MCAT-deficient T cells compared to controls (**Fig. 5B-C**, **Fig. S9B-C**). To test whether iron mis-handling resulted in increased ferroptosis, which might contribute to phenotypes of T cell dysfunction, we assessed lipid peroxidation via BODIPY C11 staining, but found no difference between M-KO CD8^+^ T cells and controls (**Fig. S9D**). We also examined ferroptosis regulators in our RNA-seq dataset, but found no consistent signature that would indicate altered susceptibility to ferroptosis in M-KO T cells relative to controls (**Fig. S9E**). We therefore concluded that elevated mitochondrial iron levels were unlikely to be acting via regulation of ferroptosis in M-KO T cells.

**Figure 5:**
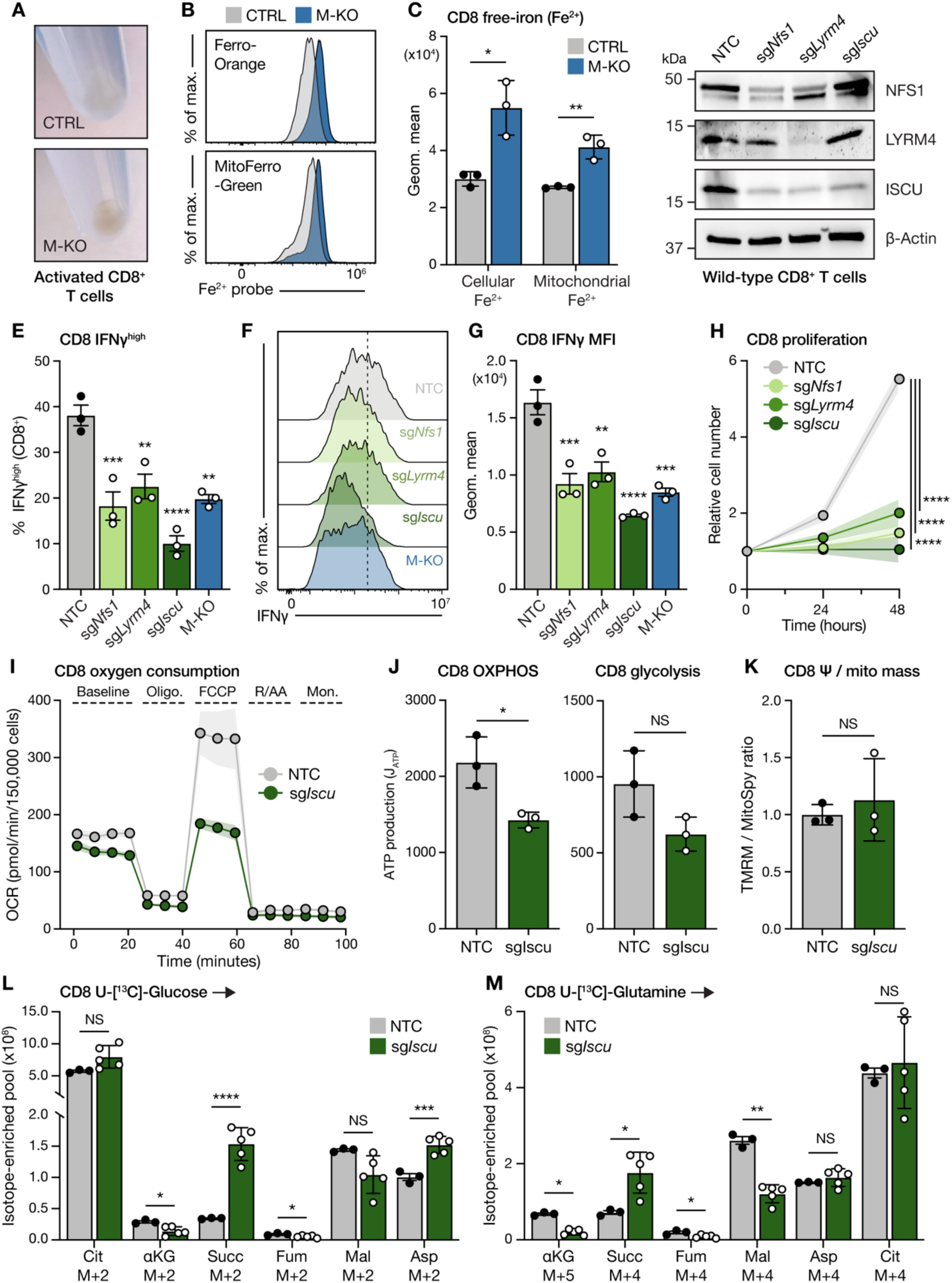
mtFAS supports effector T cell function via regulation of ISC synthesis. (**A**) Representative pictures of the pellets from activated CTRL and M-KO CD8^+^ T cell cultures. (**B-C**) Representative flow cytometry histograms (**B**) and quantification of cellular (Ferro-Orange) and mitochondrial (MitoFerro-Green) free iron (Fe^2+^) levels in CTRL and M-KO CD8^+^ T cells (**C**). Cells were activated in vitro using plate-bound anti-CD3 and anti-CD28 antibodies for 3d; data represent the mean ± SEM (*n* = 3 mice per group). (**D**) Immunoblot of NFS1, LYRM4, ISCU and β-Actin protein levels in wild-type (NTC), *Nfs1*-deleted (sg*Nfs1*), *Lyrm4*-deleted (sg*Lyrm4*), and *Iscu*-deleted (sg*Iscu*) CD8^+^ T cells. NTC denotes non-targeting control. (**E-G**) Cytokine production by wild-type CD8^+^ T cells with disrupted ISC production (sg*Nfs1*, sg*Lyrm4*, sg*Iscu*) following a 4h PMA/ionomycin stimulation. Shown are the percentage of IFN-ɣ^high^ CD8^+^ T cells (**E**), representative histograms of IFN-ɣ levels (**F**), and the mean fluorescence intensity (MFI) of IFN-ɣ expression (**G**) for each CD8^+^ T cell population shown. IFN-ɣ production in M-KO CD8^+^ T cells is shown in parallel. Data represent the mean ± SEM (*n* = 3 mice per group), significance is shown relative to the NTC group. (**H**) Proliferation of activated wild-type (NTC), *Nfs1*-deleted, *Lyrm4*-deleted, and *Iscu*-deleted CD8^+^ T cells cultured with IL-2 (50 U/mL). Cells were activated for 3d prior to CRISPR-mediated genetic deletion and allowed to recover for 2d prior to assay seeding. The number of viable T cells was then counted over the following 2d. Data represent the mean ± SEM (*n* = 3 mice per group). (**I**) Oxygen consumption rate (OCR) of activated wild-type (NTC) and *Iscu*-deleted CD8*^+^* T cells, as quantified using the Seahorse mitochondrial stress test. Data show one representative technical replicate (*n* = 3 technical replicates per group using cells pooled from 3 mice), error bars denote SD. (**J**) Calculated maximal ATP production rates derived from oxidative phosphorylation (OXPHOS) and glycolysis in wild-type (NTC) and *Iscu*-deleted CD8^+^ T cells. Data represent the mean ± SD (*n* = 3 technical replicates per group using cells pooled from 3 mice). (**K**) Quantification of mitochondrial membrane potential (Ψ) normalized to cellular mitochondrial content in wild-type (NTC) and *Iscu*-deleted CD8^+^ T cells. Data represent the mean ± SD (*n* = 3 technical replicates per group using cells pooled from 3 mice), see also Figure S9G for individual MitoSpy and TMRM stainings. (**L-M**) Contribution of [U-^13^C]-glucose (**L**) and -glutamine (**M**) to TCA cycle intermediates and aspartate in wild-type (NTC) and *Iscu*-deleted CD8^+^ T cells. Data shown represent the total abundance of the indicated metabolite mass isotopologues derived from the labeled tracer, see also Figure S9H-J for fractional enrichment data. Error bars denote SD (*n* = 3 NTC and 5 sg*Iscu* technical replicates using cells pooled from 3 mice). For statistics: \**p* > 0.05, \*\**p* < 0.01, \*\*\**p* < 0.001, \*\*\*\**p* < 0.0001 by unpaired *t* test with Welch’s correction (**C, J and K**), one-way ANOVA with Tukey’s multiple comparisons test (**E, G, L and M**), NS denotes not significant.

Beyond ferroptosis, the finding that mitochondrial iron levels were elevated in M-KO cells was consistent with the idea that these cells display reduced incorporation of iron into the ISC biosynthesis pathway, which occurs primarily in the mitochondria. To test whether the core ISC complex is needed for T cell effector function, we disrupted ISC production in wild-type CD8^+^ T cells via CRISPR/Cas9 gene editing using single-guide RNA (sgRNA) sequences targeting *Nfs1*, *Lyrm4* and *Iscu* (**Fig. 5D**). Both IFN-ɣ production (**Fig. 5E-G**) and proliferation (**Fig. 5H**) of CD8^+^ T cells were significantly reduced upon *Lyrm4*, *Nfs1*, or *Iscu* deletion compared to T cells edited with a non-targeting control (NTC) sequence. Thus, deletion of the ISC production complex mimics the reduced proliferation and effector function of *Mcat*-deficient CD8^+^ T cells.

To further explore this finding, we assessed the bioenergetics of control (NTC) and sg*Iscu*-targeted CD8^+^ T cells by Seahorse analysis. CD8^+^ T cells with *Iscu* deletion displayed reduced basal and maximal OCR (**Fig. 5I**), indicating reduced respiratory capacity, as previously observed in M-KO cells (**Fig. 3A**). ECAR rates were also reduced in sg*Iscu* T cells compared with controls (**Fig. S9F**). Consistent with this, calculated ATP production rate from OXPHOS was significantly lower in sg*Iscu* CD8^+^ T cells compared to controls, while ATP production from glycolysis trended downward but was not statistically significant (**Fig. 5J**). Membrane potential per mitochondrial mass measurements were unchanged in sg*Iscu* cells (**Fig. 5K**, **Fig. S9G**). Similar to what we observed in M-KO CD8^+^ T cells, sg*Iscu* T cells displayed ^13^C-labeling patterns consistent with decreased SDH activity, marked by a significant increase in the abundance of ^13^C-glucose-derived succinate (m+2), while labeling into downstream metabolites was unchanged relative to NTC cells (**Fig. 5L**, **Fig. S9H**). Similarly, when tracing with ^13^C-glutamine, sg*Iscu* CD8^+^ T cells displayed a build-up of labeled succinate, reduced labeling into malate downstream of SDH, and did not engage reductive carboxylation of glutamine (**Fig. 5M**, **Fig. S9I-J**).

Products of the mtFAS pathway separate into two “arms”: a long-chain arm, which regulates ETC assembly and ISC production, and an octanoyl-ACP arm, which feeds lipoic acid biosynthesis and TCA enzyme function^25^ (**Fig. 6A**). Ablation of ISC biogenesis can impact both ETC function and protein lipoylation as Lipoic Acid Synthase (LIAS) is itself an ISC containing protein^45^ (**Fig. 6B**). To rigorously delineate the key outputs of mtFAS activity required for T cell effector function, we targeted LIPT1, the enzyme downstream of LIAS that attaches lipoic acid to target enzymes including PDH and OGDH^46^. Deletion of *Lipt1* inhibited protein lipoylation in CD8^+^ T cells without affecting ETC complex assembly (SDHB, **Fig. 6B**). In sharp contrast to targeting *Iscu*, sg*Lipt1* cells showed no defects in cytokine production or proliferation compared with control T cells (**Fig. 6C-E**). Basal and maximal OCR did trend lower in *Lipt1*-deleted (sg*Lipt1*) CD8^+^ T cells (**Fig. 6F**), alongside an insignificant reduction in ECAR in sg*Lipt1* versus NTC CD8^+^ T cells (**Fig. S10A**). As such, both OXPHOS- and glycolysis-dependent ATP production rates in LIPT1-deleted T cells were not significantly different from controls, nor was a difference in membrane potential (per mitochondrial mass) observed (**Fig. 6F-H**, **Fig. S10B**). Similarly, we observed minimal changes in the incorporation of ^13^C-glucose and ^13^C-glutamine into TCA cycle intermediates and aspartate between sg*Lipt1* versus control CD8^+^ T cells, other than a reduction of ^13^C-labeled carbons into citrate (**Fig. S10C-G**)

**Figure 6:**
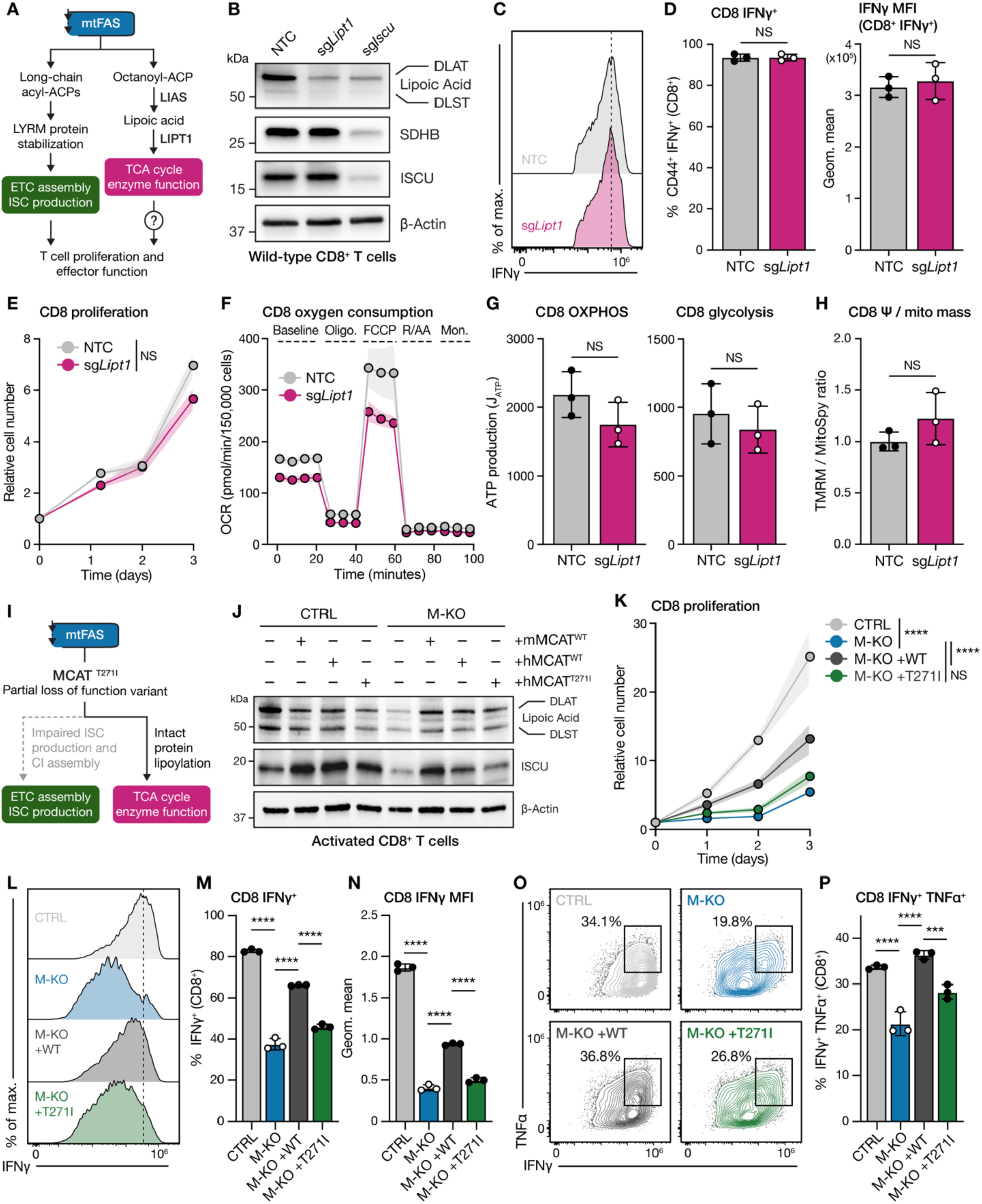
Protein lipoylation is not required for effector T cell function. (**A**) Schematic depicting how mitochondrial fatty acid synthesis (mtFAS) products support mitochondrial metabolism and effector T cell function via two distinct mechanisms. (**B**) Immunoblot of lipoylated DLAT and DLST, SDHB, ISCU and β-Actin protein levels in wild-type (NTC), *Lipt1*-deleted (sg*Lipt1*), and *Iscu*-deleted (sg*Iscu*) CD8^+^ T cells. NTC denotes non-targeting control. (**C-D**) Cytokine production by wild-type (NTC) and protein lipoylation-deficient (sg*Lipt1*) CD8^+^ T cells, following a 4h PMA/ionomycin stimulation. Shown are representative histograms of IFN-ɣ levels (**C**), percentage of CD44^+^ IFN-ɣ^+^ CD8^+^ T cells and mean fluorescence intensity (MFI) of IFN-ɣ expression (**D**). Data represent the mean ± SD (*n* = 3 technical repeats using cells pooled from 3 mice). (**E**) Proliferation of activated wild-type (NTC) and *Lipt1*-deleted CD8+ T cells cultured with IL-2 (50 U/mL). Cells were activated for 3d prior to CRISPR-mediated genetic deletion and allowed to recover for 2d prior to assay seeding. The number of viable T cells was then counted over the following 3 days. Data represent the mean ± SD (*n* = 3 technical replicates using cells pooled from 3 mice). (**F**) Oxygen consumption rate (OCR) of activated wild-type (NTC) and *Lipt1*-deleted CD8*^+^* T cells, as quantified using the Seahorse mitochondrial stress test. Data show one representative technical replicate (*n* = 3 technical replicates per group using cells pooled from 3 mice), error bars denote SD. (**G**) Calculated maximal ATP production rates derived from oxidative phosphorylation (OXPHOS) and glycolysis in wild-type (NTC) and *Lipt1*-deleted CD8^+^ T cells. Data represent the mean ± SD (*n* = 3 technical replicates per group using cells pooled from 3 mice). (**K**) Quantification of mitochondrial membrane potential (Ψ) normalized to cellular mitochondrial content in wild-type (NTC) and *Lipt1*-deleted CD8^+^ T cells. Data represent the mean ± SD (*n* = 3 technical replicates per group using cells pooled from 3 mice), see also Figure S10B for individual MitoSpy and TMRM stainings. (**I**) Schematic highlighting the reduced function of mutant MCAT (MCAT^T271I^) which supports protein lipoylation while ETC assembly and ISC production remain impaired. (**J**) Immunoblot of lipoylated DLAT and DLST, ISCU and β-Actin protein levels in activated CTRL and M-KO CD8^+^ T cells with (+) or without (-) expression of murine wild-type MCAT (+mMCAT^WT^), human wild-type MCAT (+hMCAT^WT^) or mutant MCAT (+hMCAT^T271I^). (**K**) Proliferation of activated CTRL, M-KO, M-KO with human WT MCAT (+WT), and M-KO with mutant MCAT (+T271I) CD8^+^ T cells cultured with IL-2 (50 U/mL). Naïve cells were transduced prior to activation using plate-bound anti-CD3 and anti-CD28 antibodies, then given 3d to recover before assay seeding. The number of viable T cells was then counted over the following 3d. Data represent the mean ± SD (*n* = 3 technical replicates per group using cells pooled from 2 mice per genotype). (**L-P**) Cytokine production in CTRL, M-KO, M-KO +WT, and M-KO +T271I CD8^+^ T cells. Naïve cells were transduced before activation with plate-bound anti-CD3 and anti-CD28 for 4d, followed by a 4h PMA/ionomycin restimulation prior to analysis. Shown are representative histograms of IFN-ɣ expression (**L**), the percentage of IFN-ɣ^+^ CD8^+^ T cells (**M**), the MFI of IFN-ɣ levels in CD8^+^ T cells (**N**), representative flow plots of IFN-ɣ versus TNF-*a* expression (**O**), and the percentage IFN-ɣ^+^ TNF-*a*^+^ cells (**P**). Data represent the mean ± SD (*n* = 3 technical replicates per group using cells pooled from 2 mice per genotype). For statistics: \*\*\**p* < 0.001, \*\*\*\**p* < 0.0001 by unpaired *t* test with Welch’s correction (**D, G and H**), one-way ANOVA with Tukey’s multiple comparisons test (**M, N and P**), or two-way ANOVA with Tukey’s multiple comparisons test (**E and K**), NS denotes not significant.

As an orthogonal approach to delineate the impact of ISC biogenesis versus protein lipoylation downstream of MCAT on T cell function, we expressed a previously described patient variant of MCAT (MCAT^T271I^) in M-KO CD8^+^ T cells^47^. MCAT^T271I^ is a partial loss of function variant that retains the ability to support protein lipoylation but fails to restore ISC production and complex I assembly in the absence of WT MCAT (schematic in **Fig 6I**). Indeed, expression of mouse WT MCAT or human WT MCAT rescued ISCU levels and protein lipoylation in M-KO cells, while MCAT^T271I^ only restored protein lipoylation (**Fig. 6J**). Furthermore, in contrast to expression of human WT MCAT, expression of MCAT^T271I^ failed to efficiently rescue the proliferation or cytokine production of M-KO CD8^+^ T cells (**Fig. 6K-P**). These data, together with our results targeting *Iscu* and *Lipt1* via CRISPR/Cas9 genome editing, strongly support the conclusion that ISC biogenesis is the key output of mtFAS pathway activity that is required for CD8^+^ T cell effector function.

### Loss of mtFAS promotes CD8^+^ T cell exhaustion

One notable observation from our proteomics data was increased levels of the inhibitory receptors PD-1 and TIM-3, key markers of T cell exhaustion, in *Mcat*-deficient CD8^+^ T cells (**Fig. S11A**). Terminal exhaustion in CD8^+^ T cells is programmed by the transcription factor TOX under conditions of chronic antigen stimulation^48^. Notably, PD-1, TIM-3, and TOX protein levels were all increased in M-KO CD8^+^ T cells under acute stimulation conditions (**Fig. S11A**). A meta-analysis of RNA-seq datasets from CD8^+^ T cells isolated from LCMV-infected mice or murine hepatocellular carcinoma (HCC) tumors^49–52^ revealed that mtFAS pathway enrichment was lowest in late-stage terminally exhausted (Tex^term^) CD8^+^ T cells and highest in CD8^+^ T cells that retain their cytotoxic abilities (effector-like exhausted (Tex^eff^) T cells) (**Fig. S11B**). Next, we examined mtFAS pathway signatures in CD8^+^ tumor-infiltrating lymphocytes (TILs) sorted from murine B16 melanoma tumors and analyzed by single-cell RNA-seq (scRNA-seq)^53^. A Uniform Manifold Approximation and Projection (UMAP) of cell-specific transcripts identified 8 clusters that defined 5 major CD8^+^ T cell populations based on previously reported markers^7,54,55^: naïve-like, proliferating (effectors), effector exhausted (Tex^eff^), intermediate exhausted (Tex^int^), and terminally exhausted (Tex^term^) (**Fig. 7A**, **Fig. S11C**). Expression of *Mcat* and *Ndufab1* were highest in proliferating effector T cells (‘Prolif. 1’ cluster) and inversely correlated to the expression of *Tox* or *Pdcd1* transcripts (**Fig. 7B**). GSEA confirmed that mtFAS pathway gene expression was highest in proliferating T cell clusters and lowest in *Tox^+^* or *Pdcd1*^high^ exhausted clusters (**Fig. 7B-C**). Overall, the mtFAS pathway signature was highly expressed in effector T cells and significantly decreased in Tex^term^ cells (**Fig. 7D**), suggesting a link between mtFAS and CD8^+^ T cell functional states in vivo.

**Figure 7:**
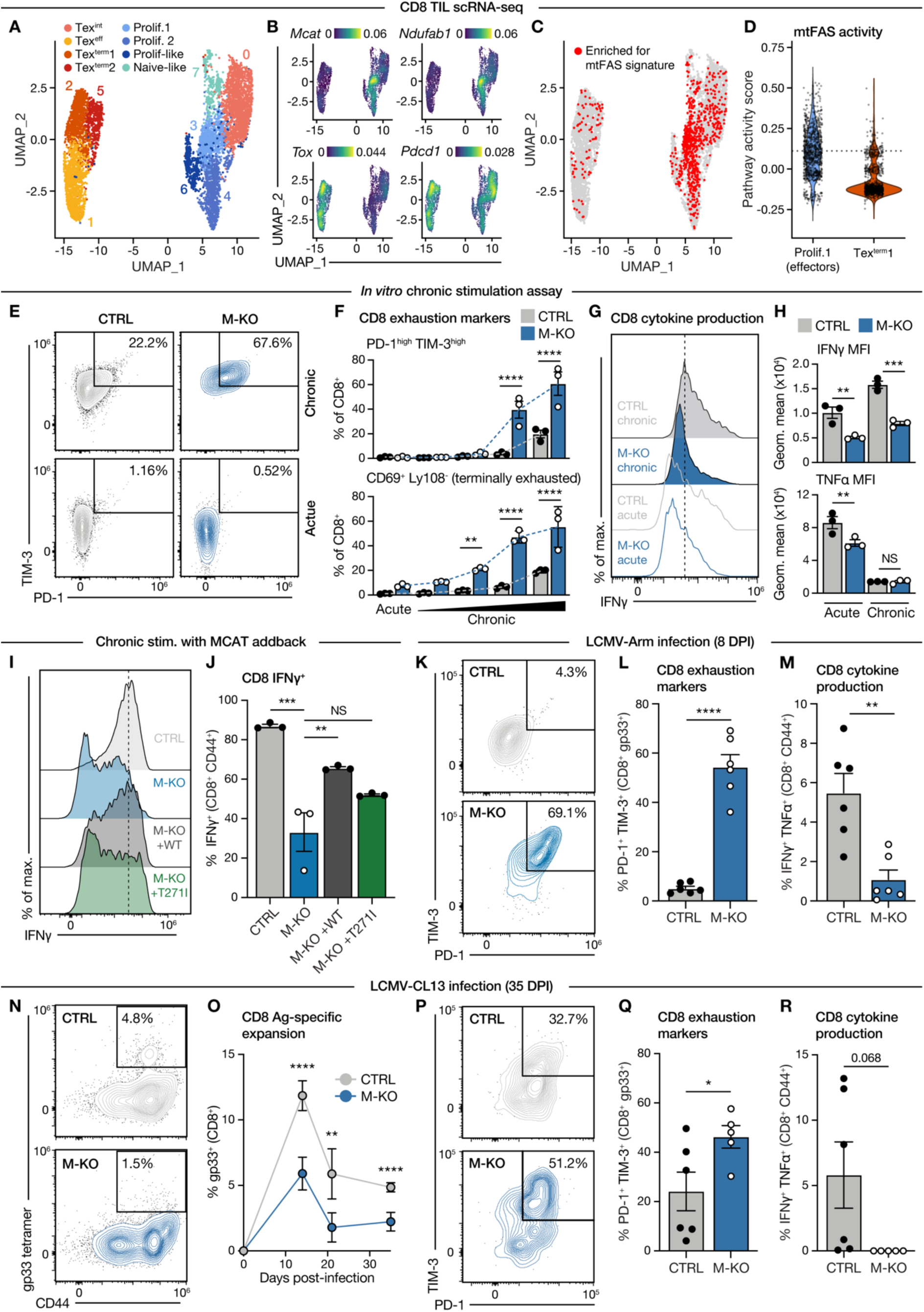
mtFAS deficiency promotes CD8^+^ T cell exhaustion. (**A-D**) Single-cell RNA sequencing (scRNA-seq) analysis of CD8^+^ tumor infiltrating lymphocytes isolated from B16 tumors. (**A**) UMAP of cell-specific transcripts identifying 8 population clusters. Effector T cell populations (*Tox*^lo^) are shown in blue and Tex cell populations (*Tox*^hi^) shown in red-yellow. (**B**) Gene expression density plots visualizing the expression of *Mcat*, *Ndufab1*, *Tox*, and *Pdcd1* overlaid on the UMAP from **A**. (**C**) GSEA embedding of mtFAS pathway gene signature enrichment overlaid on the UMAP from **A**. Colored dots represent cells with positive enrichment scores for mtFAS. (**D**) Violin plot of mtFAS pathway activity scores for Prolif.1 and Tex^term^1 clusters from **A**. See also Figure S11C for cluster identification. (**E-H**) Functional changes in CTRL and M-KO CD8^+^ T cells following chronic stimulation in vitro. Activated CTRL and M-KO CD8+ T cells were cultured under acute versus chronic *in vitro* stimulation conditions and analyzed for 3d. (**E-F**) Representative flow cytometry plots for PD-1 versus TIM-3 levels (**E**) and quantification of PD-1^hi^ TIM-3^hi^ (upper) or CD69^+^ Ly108^-^ (lower) CTRL and M-KO CD8^+^ T cells under acute and chronic *in vitro* stimulation conditions (**F**). Black slopes denote increasing Dynabead:T cell ratios (from left to right: 1:1, 2:1, 5:1, 10:1). (**G-H**) Representative flow cytometry histograms (**G**) and quantification of IFN-ɣ (upper) and TNF-*a* (lower) MFI values in CTRL and M-KO CD8^+^ T cells (**H**) under acute versus chronic in vitro stimulation conditions. Data represent the mean ± SEM (*n* = 3 mice per group). (**I-J**) Cytokine production in CTRL, M-KO, M-KO with human WT MCAT (+WT), and M-KO with mutant MCAT (+T271I) CD8^+^ T cells in response to in vitro chronic stimulation. Shown are representative histograms of IFN-ɣ expression (**I**) and quantification of IFN-ɣ^+^ CD8^+^ T cells (**J**). Data represent the mean ± SD (*n* = 3 technical replicates per group using cells pooled from 2 mice per genotype). (**K-M**) Response of M-KO CD8^+^ T cells to acute LCMV infection. CTRL and M-KO were infected with LCMV-Arm and CD8^+^ T cell responses assessed at 8 days post-infection (dpi). Representative flow cytometry plots of PD-1 versus TIM-3 levels (**K**) and quantification of PD-1^+^ TIM-3^+^ CD8^+^ gp33^+^ T cells (**L**) in the spleens of CTRL and M-KO mice at 8 dpi. (**M**) Quantification of IFN-ɣ^+^ TNF-*a*^+^ CD8^+^ CD44^+^ T cells in the spleens of CTRL and M-KO mice at 8 dpi, following *ex vivo* gp33 peptide stimulation. Data represent the ± SEM (*n* = 6 per group). (**N-R**) Response of M-KO CD8^+^ T cells to chronic LCMV infection. CTRL and M-KO mice were infected with LCMV-CL13 and CD8^+^ T cell responses assessed at 35 dpi. (**N**) Representative flow cytometry plots of gp33 tetramer^+^ versus CD44 plots for CD8^+^ T cells isolated from the spleens of CTRL and M-KO mice at 35 dpi. The percentage of gp33^+^ cells of the total CD8^+^ T cell population is shown for each genotype. (**O**) Quantification of gp33^+^ CD8^+^ T cells in the peripheral blood of CTRL and M-KO mice at 14, 21, and 35 days post-infection. Data represent the mean ± SEM (*n* = 5 mice per group). (**P-Q**) Representative flow cytometry plots of PD-1 versus TIM-3 levels (**P**) and quantification (**Q**) of PD-1^+^ TIM-3^+^ CD8^+^ gp33^+^ T cells in the spleens of CTRL and M-KO mice at 35 dpi. Data represent the mean ± SEM (*n* = 6 CTRL, 5 M-KO). (**R**) Quantification of IFN-ɣ^+^ TNF-*a*^+^ CD8^+^ CD44^+^ T cells in the spleens of CTRL and M-KO mice at 35 dpi, following *ex vivo* gp33 peptide stimulation. Data represent the mean ± SEM (*n* = 6 CTRL, 5 M-KO). For statistics: \**p* > 0.05, \*\**p* < 0.01, \*\*\**p* < 0.001, \*\*\*\**p* < 0.0001 by unpaired *t* test with Welch’s correction (**L, M, Q and R**), one-way ANOVA with Tukey’s multiple comparisons test (**J**), or two-way ANOVA with Tukey’s multiple comparisons test (**F, H and O**), NS denotes not significant.

To formally assess the role of mtFAS in T cell dysfunction, we first used an in vitro model of chronic stimulation in which CTRL and M-KO CD8^+^ T cells were subjected to either acute or chronic (repetitive) bead-based *a*CD3/CD28 stimulation^18^. By altering the ratio of *a*CD3/CD28-coated beads to T cells, we found that loss of MCAT led to a significant expansion in the number of terminally exhausted (i.e., PD-1^hi^TIM-3^hi^ or CD69^+^Ly108^-^) CD8^+^ T cells with increasing levels of chronic stimulation (**Fig. 7E-F**). Conversely, IFN-ɣ production was significantly reduced in chronically stimulated M-KO CD8^+^ T cells compared to controls (**Fig. 7G-H**). To test whether this phenotype resulted from lost protein lipoylation or ISC biogenesis and ETC function, we expressed human WT MCAT and the MCAT^T271I^ mutant that selectively rescues protein lipoylation in M-KO CD8^+^ T cells and exposed these cells to chronic stimulation with *a*CD3/CD28-coated beads. We found that expression of WT MCAT rescued cytokine production in chronically stimulated M-KO cells, while MCAT^T271I^ failed to efficiently rescue IFN-ɣ production in M-KO T cells (**Fig. 7I-J**).

Finally, we tested whether mtFAS deficiency could impact T cell exhaustion in vivo. We first infected CTRL and M-KO mice with the Armstrong strain of LCMV (LCMV-Arm), which does not normally induce T cell exhaustion in control animals^56^. However, we found that CD8^+^ T cells from infected M-KO mice displayed premature acquisition of exhaustion-associated features at only 8 days post-infection compared to control T cells, marked by increased expression of PD-1 and TIM-3 (**Fig. 7K-L**) and reduced cytokine production (**Fig. 7M**). We next infected control and M-KO mice with the Clone 13 strain of LCMV (LCMV-CL13), which establishes a chronic infection and leads to progressive CD8^+^ T cell dysfunction. Similar to our other infection models (**Fig. 2**), the expansion of LCMV-specific (gp33^+^) T cells was blunted in M-KO mice compared to controls (**Fig. 7N**), a defect that persisted over time (**Fig. 7O**). As with LCMV-Arm infection, assessment of CD8^+^ T cells in the spleen of infected animals at 35 dpi revealed a significant increase in the proportion of terminally exhausted (PD-1^+^TIM-3^+^) CD8^+^ T cells in M-KO mice compared to controls (**Fig. 7P-Q**). Similarly, this increase in inhibitory receptor expression correlated with poor effector function by *Mcat*-deficient CD8^+^ T cells, as their capacity to produce IFN-ɣ and TNF-*a* was ablated (**Fig. 7R**). Collectively, our results indicate that mtFAS is an essential mitochondrial activity required for sustaining T cell effector responses, and that loss of mtFAS activity (through targeting of MCAT) pushes CD8^+^ T cells towards dysfunction, marked by reduced effector function and acquisition of a terminal Tex phenotype.

## DISCUSSION

T cells undergo extensive mitochondrial reprogramming upon activation to support their increased metabolic demands, but the specific mitochondrial pathways that regulate T cell effector function have remained unclear. In this study, we identify mtFAS as an essential metabolic program that governs T cell function and resistance to exhaustion. By targeting the initiating enzyme MCAT, we show that loss of mtFAS activity destabilizes the core ISC biosynthesis complex (NFS1-LYRM4-ISCU). This impairs ETC complex I function and stability, reducing OXPHOS bioenergetic capacity, and resulting in sharply reduced effector cytokine production. Ultimately, these defects in mitochondrial metabolism blunt CD8^+^ and CD4^+^ T cell-mediated type I immune responses in vivo, compromising host protection against bacterial and viral infection. Moreover, MCAT loss leads to premature acquisition of exhaustion-like features in CD8^+^ T cells, highlighting a key role for mtFAS in preserving durable effector T cell responses. Together, these findings position mtFAS as a critical mitochondrial function that integrates metabolic control with maintenance of T cell effector function.

The identification of mtFAS as a key determinant of T cell effector function builds on growing recognition of the importance of this often overlooked biosynthetic pathway in cellular function and differentiation (reviewed by Wedan et al.^25^). mtFAS activity is essential in mammals, as whole-body *Mecr* deletion is embryonically lethal in mice^57^, and knocking out *Mcat* after development using a whole-body Cre results in overt metabolic insufficiencies, decreased ETC complex abundance, and shortened life span^32^. These observations have historically been attributed to the loss of TCA cycle function, owing to the pathway’s classically appreciated role in generating lipoic acid. Newer studies, however, have demonstrated that mtFAS also produces longer acyl-ACP species that engage a network of LYRM proteins to directly support ETC assembly, ISC biogenesis and mitochondrial translation^33,43,58^. Our work herein provides new understanding of the role this pathway plays in T cells: both mouse and human CD8^+^ and CD4^+^ (Th1) T cells upregulate mtFAS upon activation and require mtFAS activity to support robust effector responses and prevent terminal exhaustion.

Our findings also reveal cell type-specific and shared requirements for mtFAS in T cell effector function. While *Mcat* deletion impairs cytokine production in both CD8^+^ and CD4^+^ Th1 cells, only CD8^+^ T cells exhibit a pronounced proliferation defect. This specific effect on CD8^+^ T cell proliferation correlates with a reduction in SDH activity that is not observed in *Mcat*-deficient CD4^+^ Th1 cells. In line with the recent discovery of an alternative TCA cycle that drives changes in cell state^59^, these results may indicate that CD8^+^ T cells are more reliant on full TCA cycle turning than Th1 cells to fuel their proliferation. However, reduced TCA cycle metabolism alone cannot explain the marked loss of cytokine production observed in both M-KO CD8^+^ and CD4^+^ T cells. Instead, we identify ISC biosynthesis as the critical mtFAS output required for T cell effector function. Loss of MCAT results in the collapse of the NFS1-LYRM4-ISCU ISC biosynthetic complex, and leads to subsequent loss of ETC complex stability and function, and reduced OXPHOS bioenergetic capacity. Importantly, we demonstrate that selective deletion of ISC biosynthesis machinery (*Nfs1*, *Lyrm4*, *Iscu*) in CD8^+^ T cells phenocopies the defects in cytokine production, T cell proliferation, and bioenergetic output elicited by MCAT loss, while ablating protein lipoylation by targeting *Lipt1* does not. Moreover, selective rescue of protein lipoylation in the absence of ISC biogenesis fails to rescue phenotypes of MCAT deficiency. The data provided here provide strong mechanistic evidence that ISC biogenesis and ETC function – not protein lipoylation – are the key metabolic outputs of the mtFAS pathway that support T cell effector function.

The formation of ISCs is an essential function of mitochondria, and has been evolutionarily conserved from bacteria to humans^60^. Due to their ability to transiently bind single electrons, these clusters facilitate a variety of molecular processes, including DNA replication^61^, TCA cycle metabolism^62^, and OXPHOS (reviewed in^63^). However, while the maintenance of iron metabolism and ferroptosis have been identified as regulators of T cell proliferation, differentiation, and function^64–69^, the precise role of ISCs in T cell-mediated immune responses has been less explored. Recent work identified cysteine as a conditionally essential metabolite for the effector function of activated CD8^+^ T cells, due to the metabolism of its sulfur atom to support NFS1-dependent ISC biosynthesis^70^. Our work identifies a critical function of mtFAS in maintaining the ISC biosynthetic machinery in T cells. mtFAS sits upstream of ISC biosynthesis, as it produces the acylated ACP species required to bind and stabilize LYRM4^33,43,44^ . The loss of LYRM4, which we observed in MCAT-knockout T cells, disrupts formation of the whole ISC synthesis complex. MCAT-dependent generation of lipids attached to ACP (acyl-ACP) is therefore a key step that drives the ISC biosynthesis program in T cells. Restoring ISC production would require regeneration of this acyl-ACP product, rather than simply restoring expression of components of the biosynthesis complex. Thus, ISC biosynthesis is a critical metabolic requirement for effective CD8⁺ T cell-mediated immunity, regulated through both nutrient (cysteine) availability and mtFAS-dependent stabilization of the core ISC biosynthetic machinery.

Beyond its role in supporting T cell effector function, our work also indicates that mtFAS is a key determinant of CD8^+^ T cell function in the face of chronic antigen exposure. Across multiple models of chronic stimulation, mtFAS activity is positively correlated with T cell effector function, and low mtFAS pathway activity is associated with acquisition of a TOX-driven exhaustion program. *Mcat* deletion pushed CD8^+^ T cells more rapidly towards a terminally exhausted state, characterized by increased inhibitory receptor (i.e., PD-1, TIM-3) expression and reduced IFN-Ɣ production. These data mirror observations from other systems in which mtFAS influences lineage commitment and cell fate, such as in skeletal muscle where impairment of the pathway blocks myoblast differentiation^33^. In T cells, mitochondrial dysfunction has been previously identified as a feature of CD8^+^ T cell exhaustion^18,20,71,72^, but clear evidence of causality has been lacking. Recent work has dissected the individual roles of specific ETC complexes in determining T cell fate: complex I and complex II integrate with distinct transcriptional and metabolic programs to support CD4^+^ Th1 cell activation and effector function^22^, and complex III-derived reactive oxygen species are necessary for the development of memory CD8^+^ T cells^21^. Our data indicate that mtFAS is required for these events. mtFAS activity, via MCAT, directly controls ISC production, thereby enabling ETC complex function and providing metabolic support for T cell effector cytokine production and resistance to exhaustion. Previous work has linked MECR, another enzyme in the mtFAS pathway, to pathogenic CD4^+^ T cell function in autoimmune models^73^. Thus, the mtFAS pathway may offer multiple therapeutic targets to modulate T cell responses by manipulating mitochondrial metabolism.

## RESOURCE AVAILABILITY

### Lead Contact

Additional information and requests for raw data or reagents should be directed to, and will be made available by, the corresponding authors, Drs. Sara M. Nowinski and Russell G. Jones.

### Materials availability

All unique, stable reagents generated in this study will be made available by the Lead Contact (Dr. Sara M. Nowinski) and with a completed Material Transfer Agreement. Plasmids generated in this study will be deposited to Addgene.

### Data and code availability

Data are available in the main text, figures, supplemental files. Proteomics data are available upon request and will be deposited to the ProteomeXchange Consortium via the PRIDE partner repository prior to publication. RNAseq data are available upon request and will be deposited to the NCBI GEO repository prior to publication. scRNA-seq analysis pipelines will be uploaded to GitHub prior to publication.

## ACKNOWLEDGMENTS

We thank members of the Burton, Jones, Krawczyk, Lien, and Nowinski laboratories for scientific discussions contributing to this manuscript. We thank Jeanie Wedberg and Margene Brewer for administrative assistance. We thank members of the Van Andel Institute (VAI) Core Facilities for technical assistance, including Mass Spectrometry (RRID: SCR_024903), Flow Cytometry (RRID: SCR_022685), Bioinformatics and Biostatistics (RRID: SCR_024762), Vivarium (RRID: SCR_023211). We especially thank Christine Isaguirre, Molly Hopper, H.J. Lee, and Colt Caplan (Mass Spectrometry), Brandon Bonnema and Emma Woodcock (Vivarium), and Rachael Sheridan (Flow Cytometry) for their technical support. S.E.J.P. is supported by a VAI Metabolism & Nutrition (MeNu) Program (RRID: SCR_027494) Pathway-to-Independence Award. M.J.W. is supported by a Damon Runyon Fellowship Award (DRG 2495-23). M.S.D. is supported by a CIHR Fellowship (MFE-403514), and a National Institute of Health (NIH) Pathway to Independence K99 Award (1K99CA293261-01A1). R.G.J. is supported by the Paul G. Allen Frontiers Group Distinguished Investigator Program, Chan Zuckerberg Initiative (CZI), National Cancer Institute (NCI), National Institutes of Allergy and Infectious Diseases (NIAID), and VAI. S.M.N. is supported by the National Institute for General Medical Sciences (NIGMS), the Eunice Kennedy Shriver National Institute of Child Health and Human Development (NICHD), and the National Institute of Arthritis, Musculoskeletal, and Skin diseases (NIAMS). This work was supported by funds to R.G.J. and S.M.N. from VAI and to S.M.N. from NIGMS (R35GM151245).

## AUTHOR CONTRIBUTIONS

Conceptualization, S.E.J.P., M.J.W., R.G.J., and S.M.N.; Experimental Design, S.E.J.P., M.J.W., M.S.D., J.L., R.D.S., R.G.J., and S.M.N.; Investigation, S.E.J.P., M.J.W., L.M.D., M.S.D., J.L., I.P., S.M.KG., M.C., P.R.N., N.G., B.M.O., N.M.F.; Data Analysis, S.E.J.P., M.J.W., M.S.D., J.L., B.M.O., I.P., R.D.S., R.G.J., and S.M.N.; Writing, S.E.J.P, R.G.J., and S.M.N.; Visualization, S.E.J.P.; Supervision, R.D.S., R.G.J., and S.M.N.; Funding Acquisition, R.G.J. and S.M.N.

## MATERIALS AND METHODS

### Mouse models

All experiments were performed using either male or female C57BL6/J background mice aged 8-14 weeks. Mice were maintained under specific pathogen-free condition at the VAI. Mice were house under 12-hr light:dark cycle with continuous access to water and standard 5010 chow provided freely (LabDiet). Genotyping was performed using DNA extracted from tail or ear biopsies. The following mouse strains were from The Jackson Lab (JAX) and used in this study: C57BL/6, B6.PL-*Thy1^a^*/CyJ (CD90.1), B6.SJL-*Ptprc^a^ Pepc^b^*/BoyJ (CD45.1^+^), Tg(TcraTcrb)1100Mjb (OT-I) and B6.Cg-Tg(Cd4-Cre)1Cwi/BfluJ (*Cd4-Cre*). *Mcat*^fl/fl^ mice were originally generated by Smith et al. and kindly donated by Dr. Jianping Ye (Pennington Biomedical Research Center, LSU). B6.Gt(ROSA)26Sor^tm1(NDI1)Ncdl^ (LSL-NDI1) mice were originally generated by McElroy et al. and kindly donated by Dr. Navdeep Chandel (Northwestern Medicine). Under the VAI Institutional Animal Care and Use Committee (IACUC)-approved protocols, *Mcat*^fl/fl^ mice were crossed to *Cd4-Cre* mice, followed by subsequent breeding of Mcat^fl/+^ heterozygotes to generate a stable *Cd4-Cre;Mcat*^fl/fl^ mouse colony.

*Mcat^fl^*^/fl^;*Cre*^+^ (MCAT-knockout, M-KO) mice were matched with *Mcat^fl^*^/fl^;*Cre*^-^ (control, CTRL) littermates for use in experiments. *Cd4-Cre;Mcat*^fl/fl^ mice were crossed to P14, OT-I, or LSL-NDI1 mice using a similar breeding scheme to generate *Cd4-Cre*; *Mcat^fl/fl^* P14, *Cd4-Cre*; *Mcat^fl/fl^* OT-I, or *Cd4-Cre*; Mcat^fl/fl^ LSL-NDI1 colonies. Mouse husbandry was facilitated by the VAI Vivarium (RRID:SCR_023211).

### Primary T cell isolation and culture

Total CD8^+^, total CD4^+^, or naïve CD4^+^ T cells were purified from mouse spleens through negative selection using magnetic bead-based isolation kits (STEMCELL Technologies). Cells were cultured in Iscove’s Modified Dulbecco’s Medium (IMDM; Wisent 319-105-CL) supplemented with 10% Nu-Serum IV culture supplement (Corning), 50 U/mL penicillin and 50 μg/mL streptomycin (Gibco), 50 μM 2-mercaptoethanol (Gibco), and L-glutamine (Gibco; 6 mM final concentration). Isolated T cells were activated in vitro via stimulation with plate-bound anti anti-CD3ε (clone 145-2C11; 2 μg/mL) and anti-CD28 (clone 37.51; 1 μg/mL) antibodies for 48 hr at a cell density of 1x10^6^ cells/mL. Following activation, cells were maintained in T cell media supplemented with 50 U/mL recombinant murine IL-2 (Peprotech). Th1 cultures were also supplemented with 5 ng/mL recombinant murine IL-12 (Peprotech). Proliferating cells were re-seeded at 4x10^5^ cells/mL in fresh medium and cytokine every two days. To assess Th subtype differentiation, naïve CD4^+^ T cells were cultured for 4 days in the presence of specific cytokine cocktails. For Th1: IL-2 (10 U/mL, Peprotech), IL-12 (5 ng/mL; BioLegend). For Th17: IL-6 (20 ng/mL, Peprotech), TGF-β (2 ng/mL, BioLegend), anti-IL-4, anti IFN-ɣ and anti-IL-2 (all 2.5 μg/mL, Invitrogen), with the addition of IL-1β (20 ng/mL, Peprotech) and IL-23 (20 ng/mL, BioLegend) to induce pathogenic Th17. For Treg: IL-2 (10 U/mL, Peprotech), TGF-β (2 ng/mL, BioLegend)

### Infection with *Listeria monocytogenes* (LmOVA)

CTRL and M-KO mice were infected intravenously with recombinant attenuated *Listeria monocytogenes* expressing OVA (LmOVA) at a sublethal dose of 2x10^6^ colony-forming units (CFU). Spleens were isolated at 7 days post-infection, dissociated into a single cell suspension, and analyzed by flow cytometry. To assess antigen-specific cytokine production of CD8^+^ and CD4^+^ T cells, splenocytes were stimulated with 1 μg/mL of OVA_257-264_ (SIINFEKL) or LLO_190-201_ (NEKYAQAYPNVS) peptides respectively, and cultured in T cell media with GolgiStop for 4 hr. Following peptide stimulation, cells were analyzed by flow cytometry.

### Infection with lymphocytic choriomeningitis virus (LCMV) and co-adoptive transfer

LCMV-Armstrong (Arm) and -Clone13 (CL13) viral particles were prepared as previously described^51^. LCMV-Arm was diluted to a concentration of 2x10^5^ plaque-forming units (PFU) per dose, and LCMV-CL13 was diluted to a concentration of 2x10^6^ PFUs per dose, both diluted in 1X PBS. CTRL and M-KO mice were infected via intraperitoneal injection (Arm) or via tail vein injection (CL13). At endpoint (8 days for LCMV-Arm, 35 days for LCMV-CL13), splenocytes were isolated from CTRL and M-KO mice stimulated with 1 μg/mL gp33 peptide (KAVYNFATM), and cultured in T cell media with GolgiStop for 4 hr. Cells were then analyzed by flow cytometry. For co-adoptive transfer experiments, 10,000 wild-type P14 CD8+ T (P14) cells (CD45.2^+^ CD90.1^+^) and 10,000 CTRL or M-KO P14 cells (CD45.2^+^ CD90.2^+^) were transferred intravenously into congenic hosts (CD45.1^+^) one day prior to infection. At 8 days post-LCMV-CL13 infection, splenocytes were isolated, stimulated and analyzed as above.

### Infection with *Citrobacter rodentium*

CTRL and M-KO mice were with *Citrobacter rodentium* DBS100 at a dose of 2-3x10^9^ colony-forming units (CFU). To monitor bacterial concentrations over time, feces were weighed and homogenized in 20% glycerol (in PBS), serially diluted, and plated on MacConkey agar (ThermoFisher) plates for calculation of CFU/mL. For immune phenotyping of the lamina propria at 7 days post-infection, the large intestine was isolated, cut longitudinally, flushed with cold HBSS buffer, and cut into small pieces. Tissue pieces were incubated in EDTA buffer (1X HBSS, 15 mM HEPES, 2% FBS, penicillin-streptomycin, 2 mM EDTA) for 15 min at 37°C. This was repeated twice, changing the buffer after each incubation. Tissue pieces were then incubated in digestion buffer [RPMI 1640 supplemented with 10 mM HEPES, 1 mM sodium pyruvate, 10% FBS, penicillin-streptomycin, 2-ME, deoxyribonuclease I (30 μg/ml; Roche), and Liberase (0.3 Wünsch U/ml; Roche)] on a magnetic stirring plate set to 600 rpm at 37°C for 50 min. The resulting cell suspensions were filtered (100 μm cell strainer) and centrifuged at 600xg for 5 min. Cells were resuspended in 40% room temperature Percoll (Cytiva) and layered onto 70% Percoll, before centrifugation at 800xg for 20 min at 20°C (no brake). Cells in the interphase were collected and analyzed by flow cytometry.

### Immunoblotting

Protein lysates were isolated from cell pellets using RIPA buffer supplemented with protease and phosphatase inhibitor cocktails (Roche) on ice for ≥30 min, vortexing frequently. Protein concentrations were quantified using a Pierce BCA Protein Assay Kit (Thermo Fisher Scientific). Equal amounts of protein were diluted in Laemmli buffer, boiled for 10 min, separated by SDS page (4-20% gradient gels), and transferred to nitrocellulose membranes. Membranes were blocked for 1 hr in 5% non-fat milk in TBST at room temperature and incubated with primary antibodies diluted 1:1000 in 5% non-fat milk overnight at 4°C (except LYRM4, diluted 1:500). Membranes were washed 3x in 1x TBST, incubated for 1 hr at room temperature a secondary antibody diluted 1:2500 in 5% milk (either HRP-conjugated, or conjugated to a DyLite 680 fluorophore). Membranes were washed 4x with 1x TBST, developed using SuperSignal West Femto Maximum Sensitivity Substrate (Thermo Scientific, cat no. PI34096), and imaged using a BioRad ChemiDoc.

### RNA isolation and qPCR analysis

Total RNA was isolated from murine T cell pellets using Qiagen’s RNeasy mini kit (cat no. 74104) following manufacturer instructions. Isolated RNA was quantified by nanodrop, reverse transcribed using a High-Capacity cDNA Reverse Transcriptase kit (Life Technologies, cat. no. 4368814), and qPCR performed using universal SYBR Green SuperMix (BioRad, cat no. 1725271). Results were normalized to *Actb* mRNA levels and control conditions using standard ΔΔCt methods. *Mcat* primers: GTCATCGACAACTGTGTTGCTGC (F), GGCTCCAGCAAATACCAAGGC (R). *Actb* primers: CATTGCTGACAGGATGCAGAAGG (F), TGCTGGAAGGTGGACAGTGAGG (R)

### Flow cytometry

Single-cell suspensions of spleens were prepared by mechanical dissociation followed by red blood cell lysis, as previously described^74^. Cell suspensions (cultured cells or splenocytes) were stained with a cocktail of fluorescently labeled antibodies and/or dyes. Cell viability was assessed using Fixable Viability Dye eFluor506 (eBioscience, cat no. 65-0866-14) according to manufacturer instructions. Cell proliferation assessed by Violet Proliferation Dye 450 (VPD450) dilution according to the manufacturer’s protocol. To assess cytokine production, cell suspensions were incubated in T cell media with 1 μg/mL peptide (OVA, LLO, or gp33), or phorbol myristate acetate (PMA; 50 ng ml^-1^) and ionomycin (500 ng ml^-1^) for 4 hr, in the presence of GolgiStop (1:1500 dilution, BD Biosciences, cat. No 554724). After stimulation, cells were stained with surface marker antibodies in staining buffer (PBS with 2% FBS and 0.02% sodium azide) at 4°C for 1 h, fixed and permeabilized at 4°C for 1 h using the Foxp3/Transcription Factor Staining Buffer Set (ThermoFisher, cat no. 00-5523-00), and then stained with intracellular marker antibodies for 1 hr. To assess free iron (Fe^2+^) levels, Ferro-Orange (Dojindo, F374) and MitoFerro-Green (Dojindo, M489) were used to stain cells according to manufacturer instructions (30 min incubation, 37°C), before immediate acquisition without fixation. Mitochondrial dyes MitoSpy-Green (2 μL, BioLegend, cat no. 424806) and TMRM (1 μL, ThermoFisher, T668) were used to stain cells according to manufacturer instructions (30 min incubation, 37°C), before immediate acquisition without fixation. To quantify fatty acid uptake, BODIPY FL C_12_ (Invitrogen, D3822) was used to stain cells according to manufacturer instructions (1 μM, 25 min incubation, 37°C). To assess lipid peroxidation, BODIPY 581/591 C11 (Invitrogen, D3861) was used to stain cells according to manufacturer instructions (2 μM, 25 min incubation, 37°C). Flow cytometry was performed on an Aurora Cytek cytometer. Analysis of .fcs files was performed using FlowJo software (TreeStar). Flow cytometry experiments were supported by the VAI Flow Cytometry Core (RRID:SCR_022685).

### Extracellular flux analysis (Seahorse)

T cell oxygen consumption rate (OCR) and extracellular acidification rate (ECAR) were measured by a Seahorse XF96 Extracellular Flux Analyzer as previously described^75^. Briefly, 150,000 activated CD8^+^ or Th1 cells were seeded in Seahorse XF medium containing 5 mM glucose, 0.5 mM glutamine, and 1 mM sodium pyruvate. Cells were centrifuged in a poly-D-lysine coated XF96 plate to ensure adherence. Cellular bioenergetics were assessed at regular time intervals according to a mito stress test protocol: sequential addition of oligomycin (2 μM), carbonyl cyanide 4-(trifluoromethoxy)phenylhydrazone (FCCP; 1.5 μM), rotenone/antimycin A (0.5 μM each), and monensin (20 μM). Data were normalized to mitochondrial content (see Flow Cytometry section for MitoSpy staining) for OCR analysis, and normalized to cell number for ECAR analysis. ATP production values were calculated based on protocols developed by Mookerjee *et al*. ^76^, as previously described.

### ^13^C stable isotope labeling

CTRL and M-KO CD8^+^ and Th1 cells were activated in vitro as described above and expanded for 3 days in T cell media. Cells were washed with VIM medium^77^ and seeded in a 24-well plate (2x10^6^ cells per well) for 4 hr in VIM containing 5 mM [U-^13^C]-glucose or 0.5 mM [U-^13^C]-glutamine, 10% dialyzed FBS (dFBS), and 50 μM 2-mercaptoethanol, and supplemented with recombinant IL-2 (and IL-12 for Th1 culture). After labeling, cells were pelleted in 1.5 mL Eppendorf tubes (600xg, 2 min, 4°C) and washed once with ice-cold saline before being snap-frozen on dry ice and stored at -80°C. Metabolomics was performed as previously described^78^ by the VAI Mass Spectrometry Core (RRID:SCR_024903).

### Proteomics and gene set enrichment analysis (GSEA)

2-3x10^6^ CTRL and M-KO CD8^+^ and Th1 cells, activated and expanded in vitro, were pelleted in 1.5 mL Eppendorf tubes, washed twice with ice-cold saline, and snap-frozen on ice before storage at -80°C. Cell lysates were extracted and digested using EasyPrep MS Sample Prep Kits (Thermo Fisher Scientific) according to the manufacturer’s instructions. Dried samples were resuspended in 0.1% trifluoroacetic acid. Global proteome quantitation was performed as previously described ^79^by the VAI Mass Spectrometry Core (RRID:SCR_024903). The resulting proteomics data were statistically tested by the VAI Bioinformatics and Biostatistics Core (RRID:SCR_024762). Briefly, differential abundance analysis of 7985 proteins was performed in R (v 4.5.0). All proteins with 0 variance, or more than 25% missingness across samples, were removed from analysis. (n = 1). Proteins with more than 25% missingness across samples were removed from analysis (n = 210). The remaining proteins (n = 7776) were log2 transformed and imputed before running limma (under default settings) and eBayes (using both trend and robust options). All p-values were adjusted together using the Benjamini-Hochberg method. Functional enrichment analyses were performed on preranked fold change data using GSEA soft-ware (Broad Institute, version 4.1.0). GSEAPreranked was run using default settings against the KEGG molecular signature database C2 and C5 on the “Mouse Gene Symbol Remapping Human Orthologs MSigDB.v.2023.2.Hs.chip” platform.

### RNA sequencing (RNA-seq) and GSEA

Approximately 0.5x10^6^ cells were collected by centrifugation, washed once with cold PBS, and resuspended in 55 μL DNA/RNA shield (Zymo Research, R1100-50). Samples were submitted to Plasmidsaurus for 3’ end counting RNA-seq. Briefly, library preparation used poly(dT)VN reverse transcription with unique molecular identifier tagging. Sequencing was performed on an Illumina NovaSeq X Plus platform generating approximately 90 bp asymmetric paired-end reads. Demultiplexing, quality filtering (fastp), alignment (STAR), UMI deduplication (UMICollapse), and gene-level quantification (featureCounts) were conducted via Plasmidsaurus’ automated bioinformatics pipeline. GSEAPreranked was performed using the same protocol as for proteomics data (see above) against the MH Hallmarks signature database.

### Rescue of MCAT expression

Naïve CD8^+^ T cells were isolated as described above and infected with retroviruses expressing the murine wild-type MCAT CDS, human wild-type MCAT, human MCAT harboring the T271I mutation^47^, or an empty vector control. Wild-type MCAT sequences were taken from the NCBI RefSeq database (NM_001030014.3 and NM_173467.5), with the T271I point mutant engineered from the human sequence. MSCV retroviral expression constructs were generated by Vector Builder using provided MCAT sequences. Virus was packaged in HEK293T cells, collected at 48-72 hr post-transfection, and viral particles were concentrated by centrifugation following incubation with Lenti-X concentrator (Takara Bio, 16 hr incubation at 4°C). Concentrated virus was supplemented with 8 μg/mL polybrene and then used to infect naïve T cells via spin transduction (1180g, 30°C, 90 min). Post-transduction, T cells were immediately activated via plate-bound anti-CD3/CD28, as outlined above. A media change was performed after approximately 24 hr, and cells were allowed to activate and proliferate for a further 48 hr prior to use in subsequent assays.

### CRISPR/Cas9-mediated gene editing

Electroporation of primary CD8^+^ T cells was adapted from the method developed by Nussing *et al*.^80^. First, CD8^+^ T cells were isolated from wild-type C57BL/6 mice according to the methodology outlined above. A single guide RNA (sgRNA) targeting murine *Nfs1*, *Lyrm4*, *Iscu, Lipt1*, as well as a non-targeting control (NTC) sgRNA were obtained from Synthego (CRISPRevolution sgRNA EZ kit). 1.86 μL of *Sp*-Cas9 nuclease (Synthego) was combined with 3.4 μL of target sgRNA (0.1 nmol/μL stock in nuclease-free water) to form sgRNA/Cas9 ribonucleoprotein (RNP) complexes, and incubated for 10 min at room temperature. While incubating, 2.5x10^6^ CD8^+^ T cells were pelleted, washed twice in PBS, and resuspended in 20 μL of P3 buffer (P3 Primary Cell 4D-Nucleofector X Kit S, Lonza). Cells were added to the RNP complex mix, before transferring the total volume to one well of a 16-well Nucleocuvette Strip (Lonza). Cells were electroporated using program DN 100, followed by immediate addition of 150 μL of pre-warmed T cell media. Cells were then transferred to 12-well plats and expanded for experimental use.

### RNAseq meta-analysis and single-sample GSEA (ssGSEA)

We mined the RNAseq meta-analysis dataset generated by Dahabieh *et al*.^51^ from previously-published data (GEO accession numbers: GSE89307, GSE84820, GSE86881), focusing on the sequencing data from CD8^+^ T cells isolated from LCMV-infected mice and mice bearing liver-tumors. ssGSEA was subsequently performed according to methodology described in Barbie *et al*.^81^, using a self-defined mtFAS pathway gene set *(Mcat, Oxsm, Cbr4, Hsd17b8, Htd2, Mecr, Ndufab1)* and gene sets that define effector exhausted (Tex^eff^) and terminally exhausted (Tex^term^) CD8^+^ T cell populations, as previously described by Daniel *et al*. ^7^. Analysis was performed on the GenePattern cloud–based public server, using the “ssGSEA” mod-ule (v10.1.0). RNA expression Z-scores were used as input data, and default settings were used for all other parameters.

### In vitro chronic stimulation assay (T cell exhaustion model)

Acute versus chronic T cell stimulation experiments were conducted in a similar fashion to previously described protocol^18^. Briefly, CTRL and M-KO CD8^+^ T cells were isolated, activated via plate-bound anti-CD3 and anti-CD28 stimulation, and expanded in T cell media supplemented with IL-2, as detailed above. At 48 hr post-activation, cells were seeded into a round-bottom 96-well plate (20,000 cells per well) under different stimulatory conditions. For chronic stimulation, T cells were incubated with increasing concentrations of CD3/CD28 Dynabeads (ThermoFisher, 11453D); tested bead:cell ratios were 1:1, 2:1, 5:1, and 10:1. Acute stimulation samples were cultured at the same cell density, but without the presence of Dynabeads. T cells were analyzed by flow cytometry after 48 hr of stimulation under these conditions (4 days total post-activation).

### CD8^+^ TIL scRNAseq analysis

Single cell (sc)RNAseq analysis of CD8^+^ T cell populations from B16 tumors was conducted as previously described^53^ on previously published data (GEO accession: GSE267070). To generate a joint uniform manifold approximation and projection (UMAP) embedding that integrates both GEX and ADT data, the FindMultiModalNeighbors function was used, followed by the RunUMAP function, resulting in a weighted nearest-neighbor UMAP. Cluster identification was achieved by integrating gene expression profiles, ADT data and gene expression density visualizations using the plot_density() function from the Nebulosa package (v.1.14.0). Details of cluster identification are provided in Figure S6. Statistical analyses for violin plots were performed using the Kruskal–Wallis test, followed by pairwise Wilcoxon comparison tests with Bonferroni *P* value adjustment.

### Statistical analysis

Prism 10 software (GraphPad) was used to perform statistical testing, unless stated otherwise. *P* values for the differences of means between groups were calculated using unpaired *t* tests with Welch’s correction, one-way ANOVA and two-way ANOVA with multiple comparison tests as appropriate, or Pearson rank-order correlations. *P* values less than 0.05 were considered statistically significant.

