## Supplemental Figures for "Mitochondrial fatty acid synthesis (mtFAS) fuels iron-sulfur cluster production essential for type I immunity"

Supplemental Figures (Preston et al.)

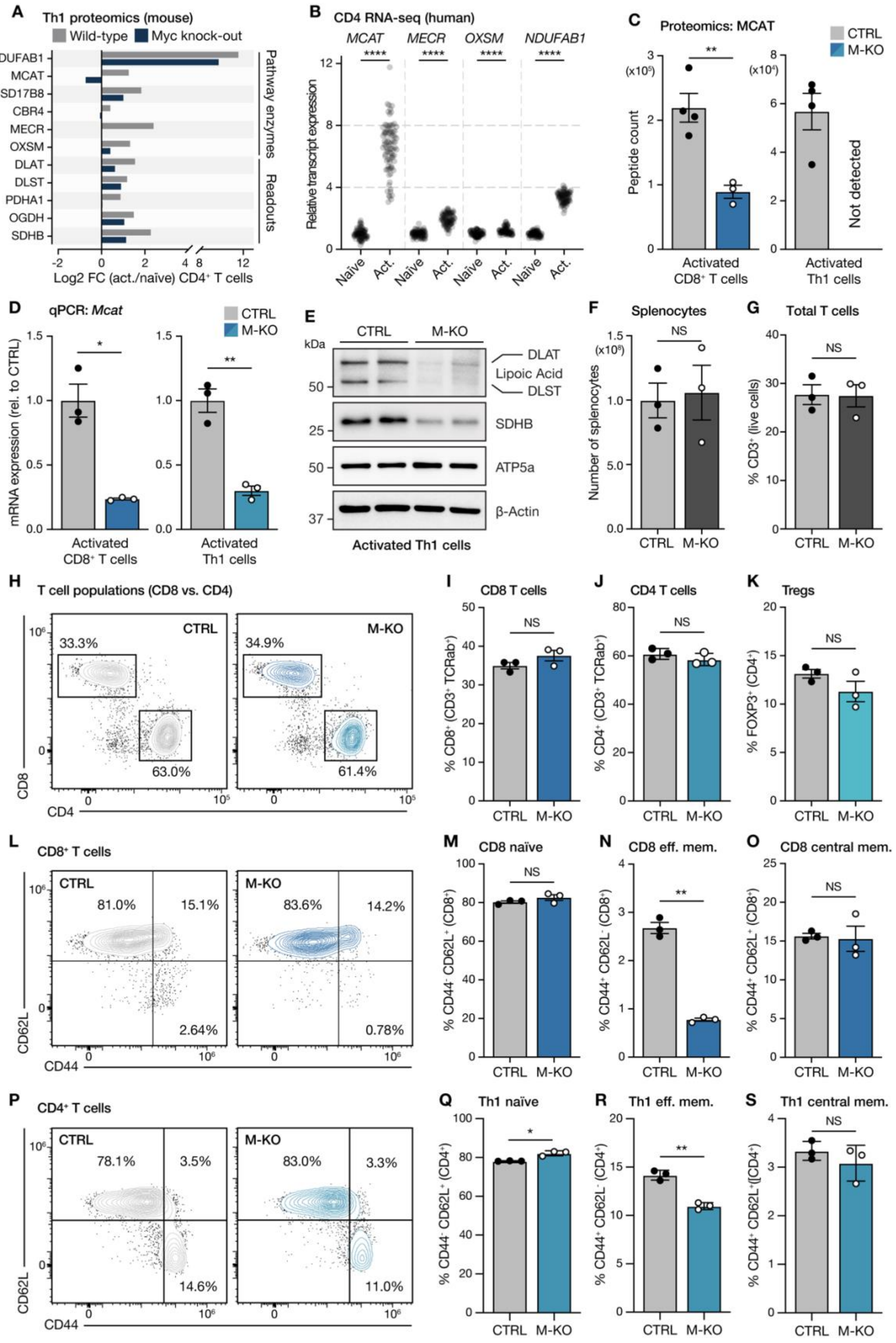

**Figure S1, related to Figure 1: Characterization of T cell *Mcat*-deficient mice.** (**A**) Abundance of mtFAS pathway components and downstream mtFAS-dependent proteins in activated $CD4^+$ Th1 cells from wild-type and Myc knockout ($Myc^{fl/fl}$;$Cd4Cre^+$) mice. Data are from the ImmPRes Myc-regulated T cell proteome dataset and displayed as Log2 fold change in activated cells relative to naïve. (**B**) RNA expression of mtFAS pathway genes in human naïve and activated $CD4^+$ T cells isolated from peripheral blood. Data are from the DICE database and shown as transcripts per million. (**C**) MCAT peptide counts in activated CTRL and M-KO $CD8^+$ T (left) and $CD4^+$ Th1 (right) cells. Data represent the mean ± SEM ($n$ = 4 CTRL, 3 M-KO). (**D**) Relative expression of *Mcat* mRNA in activated CTRL and M-KO $CD8^+$ T (left) and $CD4^+$ Th1 (right) cells as measured by qPCR. Data represent the mean ± SEM ($n$ = 3 mice per group). (**E**) Immunoblot of lipoylated DLAT and DLST, SDHB, ATP5a and β-Actin in activated CTRL and M-KO $CD4^+$ Th1 cells. (**F**) Total number of splenocytes in CTRL and M-KO mice at 8-12 weeks of age. Data represent the mean ± SEM ($n$ = 3 mice per group). (**G**) Percentage of $CD3^+$ cells in the spleens of CTRL and M-KO mice at 8-12 weeks of age. Data represent the mean ± SEM ($n$ = 3 mice per group). (**H-K**) Representative flow cytometry plots of CD8 versus CD4 expression for $CD3^+$ splenocytes (**E**) and quantification of the percentage of $CD8^+$ (**F**), $CD4^+$ (**G**), or $FOXP3^+$ (**H**) cells in the spleens of CTRL and M-KO mice under baseline conditions. Data represent the mean ± SEM ($n$ = 3 mice per group). (**L-O**) Representative flow cytometry plots of CD44 versus CD62L (**L**) and quantification of the percentage $CD44^-$ $CD62L^-$ (**M**), $CD44^+$ $CD62L^-$ (**N**), or $CD44^+$ $CD62L^+$ (**O**) $CD8^+$ T cells in the spleens of CTRL and M-KO mice under baseline conditions. Data represent the mean ± SEM ($n$ = 3 mice per group). (**P-S**) Representative flow cytometry plots of CD44 versus CD62L (**P**) and quantification of the percentage $CD44^-$ $CD62L^-$ (**Q**), $CD44^+$ $CD62L^-$ (**R**), or $CD44^+$ $CD62L^+$ (**S**) $CD4^+$ Th1 T cells in the spleens of CTRL and M-KO mice under baseline conditions. Data represent the mean ± SEM ($n$ = 3 mice per group). For statistics: *$p$ > 0.05, **$p$ < 0.01 by unpaired $t$ test with Welch's correction (**C, D, F, G, I-K, M-O and Q-S**), NS denotes not significant.

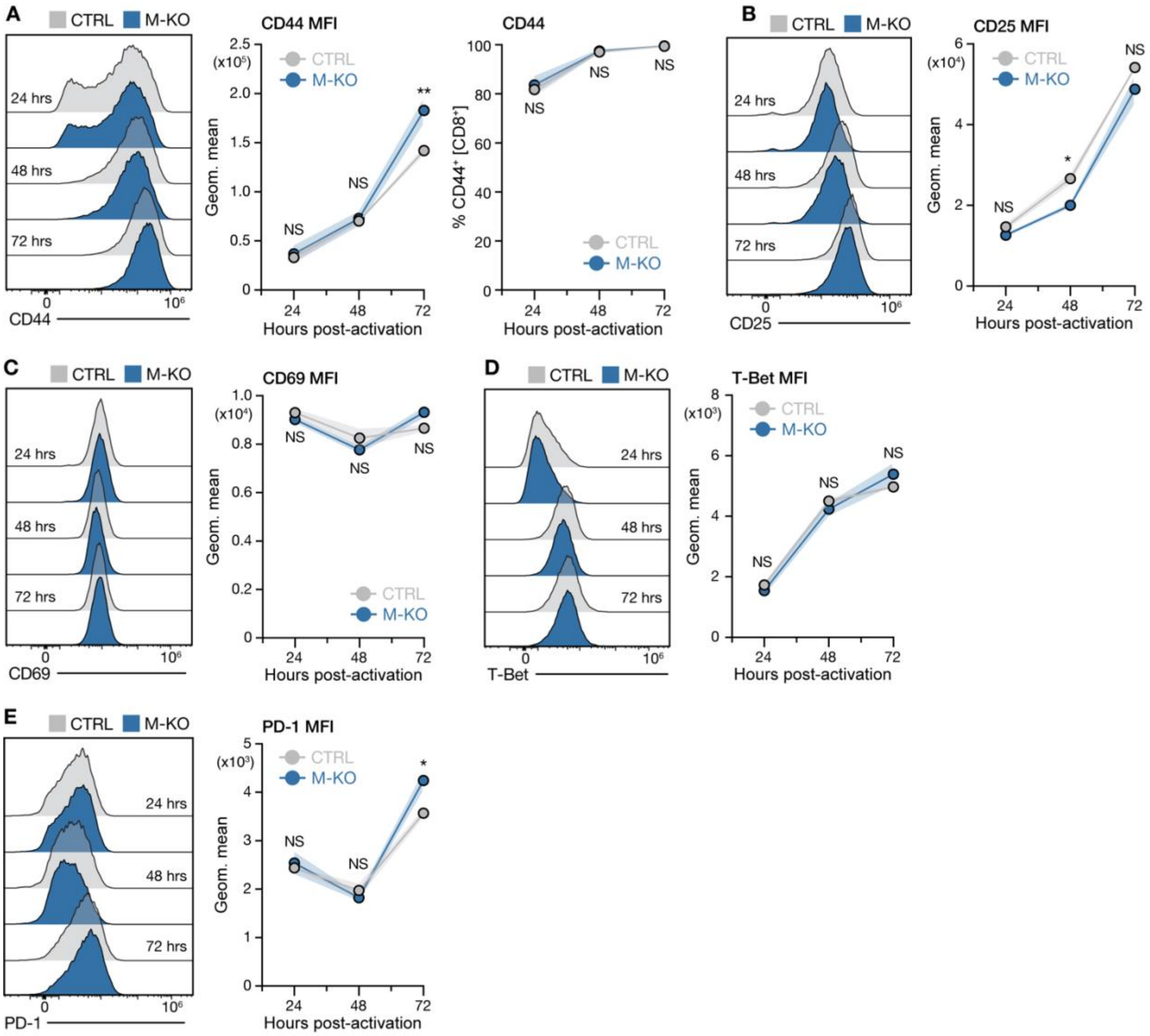


**Figure S2, related to Figure 1: CD8⁺ T cell activation dynamics are largely unchanged by MCAT loss.** Naïve CTRL and M-KO CD8⁺ T cells were activated using plate-bound anti-CD3 and anti-CD28 antibodies and the expression of CD44 (**A**), CD25 (**B**), CD69 (**C**), T-Bet (**D**), and PD-1 (**E**) were quantified every 24h for 72h total. Representative histograms are shown for the levels of each activation marker across the time-course, alongside the mean fluorescence intensity (MFI) of expression and percentage of marker-positive CD8⁺ T cells, as appropriate. Error bars denote SEM ($n$ = 3 mice per group). For statistics: *$p > 0.05$, **$p < 0.01$ by two-way ANOVA with Tukey's multiple comparisons test (**A-E**), NS denotes not significant.

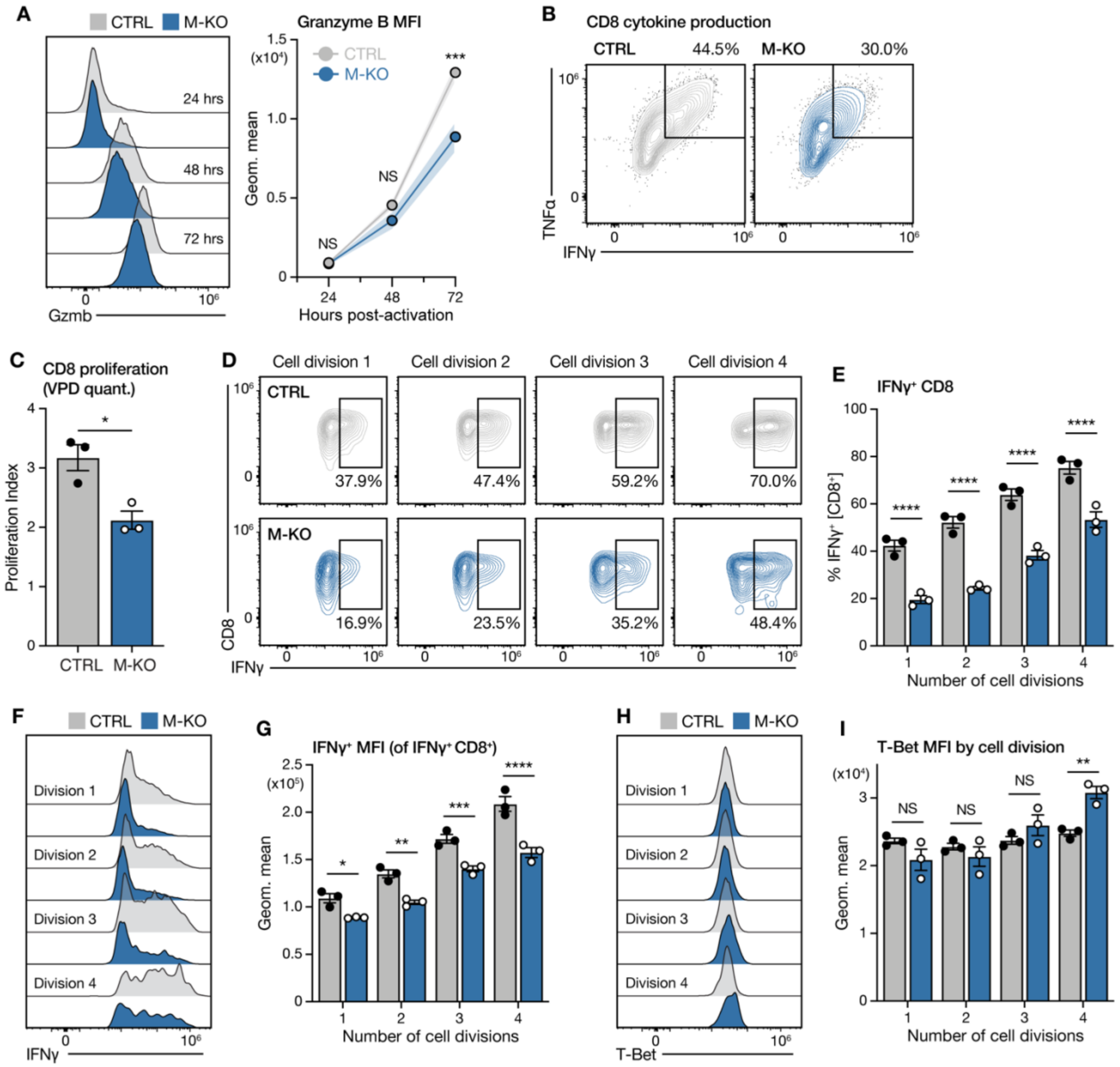


**Figure S3, related to Figure 1: Functional characterization of *Mcat*-deficient CD8$^{+}$ T cells.** (**A**) Representative histogram (left) and mean fluorescence intensity (MFI) of Granzyme B expression in CTRL and M-KO CD8$^{+}$ T cells across the first 72 hours of in vitro activation. Error bars denote SEM ($n$ = 3 mice per group). (**B**) Representative flow plot of IFN-ɣ versus TNF-α expression in activated CTRL and M-KO CD8$^{+}$ T cells, following a 4h PMA/ionomycin stimulation. The percentages of IFN-γ$^{+}$ TNF-α$^{+}$ cells are listed. (**C**) Quantification of CTRL and M-KO CD8$^{+}$ T cell proliferation index over 72 hours of cell division following in vitro activation, as measured by violet proliferation dye (VPD, see Figure 1I for representative histogram). Naïve cells were incubated with VPD for 30m prior to activation using plate-bound anti-CD3 and anti-CD28 antibodies. Data represent the mean ± SEM ($n$ = 3 mice per group). (**D**) Representative flow plots of CD8 versus IFN-ɣ expression in CTRL and M-KO CD8$^{+}$ T cells across cell divisions, as determined by VPD dilution. The percentages of IFN-γ$^{+}$ cells in each cell division are indicated. (**E-G**) IFN-ɣ production in CTRL and M-KO CD8$^{+}$ T cells across cell divisions, as determined by VPD dilution. Data shown are the percentage of IFN-γ$^{+}$ CD8$^{+}$ cells per division (**E**), representative histograms of IFN-ɣ expression per division (**F**), and the MFI of IFN-ɣ levels in IFN-ɣ$^{+}$ CD8$^{+}$ T cells per division (**G**). Data represent the mean ± SEM ($n$ = 3 mice per group). (**H-I**) Representative histograms of T-Bet expression (**H**) and MFI of T-Bet levels (**I**) in CTRL and M-KO CD8$^{+}$ T cells per cell division, as determined by VPD dilution. Data represent the mean ± SEM ($n$ = 3 mice per group). For statistics: *$p > 0.05$, **$p < 0.01$, ***$p < 0.001$, ****$p < 0.0001$ by unpaired $t$ test with Welch's correction (**C**), or two-way ANOVA with Tukey's multiple comparisons test (**A, E, G and I**), NS denotes not significant.

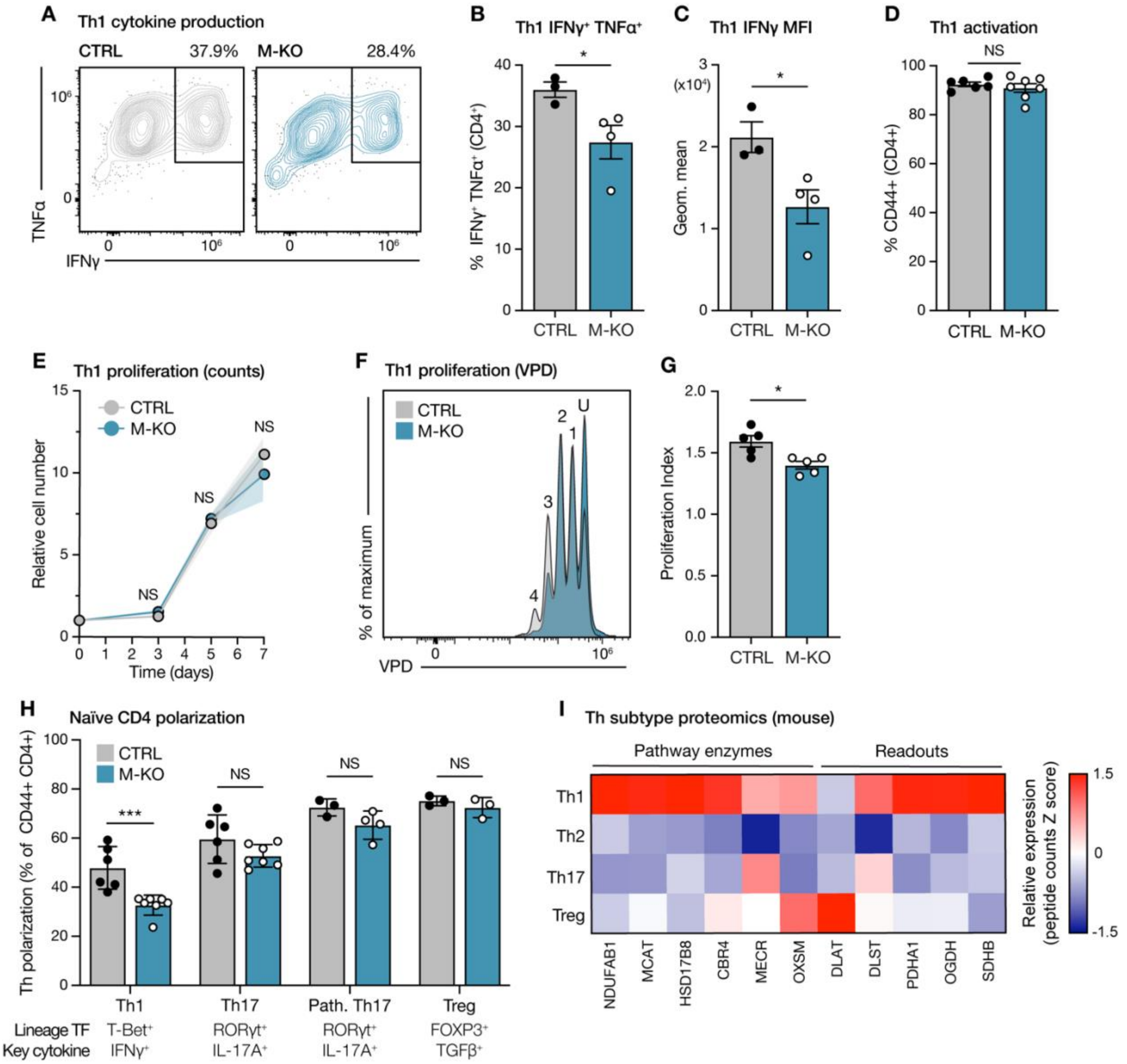


**Figure S4, related to Figure 1: Functional characterization of *Mcat*-deficient CD4⁺ Th1 cells.** (**A-C**) Cytokine production in activated CTRL and M-KO CD4⁺ Th1 cells following a 4h PMA/ionomycin stimulation. Shown are representative flow cytometry plots for IFN-ɣ versus TNF-α expression (**A**), quantification of IFN-ɣ⁺ TNF-α⁺ cells (**B**), and mean fluorescence intensity (MFI) of IFN-ɣ levels in CD4⁺ Th1 cells (**C**). Data represent the mean ± SEM ($n$ = 3 CTRL, 4 M-KO). (**D**) Percentage of CD44+ CTRL and M-KO CD4+ Th1 cells 3d post-activation with plate-bound anti-CD3 and anti-CD28. Data represent the mean ± SEM ($n$ = 6 per group). (**E-G**) Proliferation of CTRL and M-KO CD4⁺ Th1 cells cultured in IL-2 (50 U/mL) and IL-12 (5 ng/mL) following in vitro activation. Shown are the viable T cell counts over 7d of culture (E), and representative histograms of violet proliferation dye (VPD) dilution alongside quantification of proliferation index over 72h of cell division (**F**). The number of cell divisions are indicated; U denotes undivided cells. Data represent the mean ± SEM ($n$ = 5 mice per group). (**H**) Naïve CD4⁺ T cells were activated with anti-CD3 and -CD28 antibodies and polarized towards a Th1, Th17, pathogenic (path.) Th17 or Treg lineage phenotype over 4d. Shown is the percentage of CTRL and M-KO T cells (CD4$^+$ CD44$^+$) that have polarized towards each lineage. Polarization efficiency was determined by the expression of a lineage-defining transcription factor and cell type-specific cytokine, as listed. Data represent the mean ± SEM (For Th1 and Th17: *n* = 6 CTRL, 7 M-KO; for path. Th17 and Treg: *n* = 3 CTRL, 4 M-KO). (**I**) Heatmap showing relative protein levels of mtFAS pathway components and downstream readouts in Th1, Th2, Th17 and Treg cells. Data are from the ImmPRes hematopoietic cell proteome dataset and are displayed as peptide count Z scores (column-normalized). For statistics: *$p > 0.05$, ***$p < 0.001$ by unpaired t test with Welch's correction (**B-D and G**), or two-way ANOVA with Tukey's multiple comparisons test (**H**), NS denotes not significant.

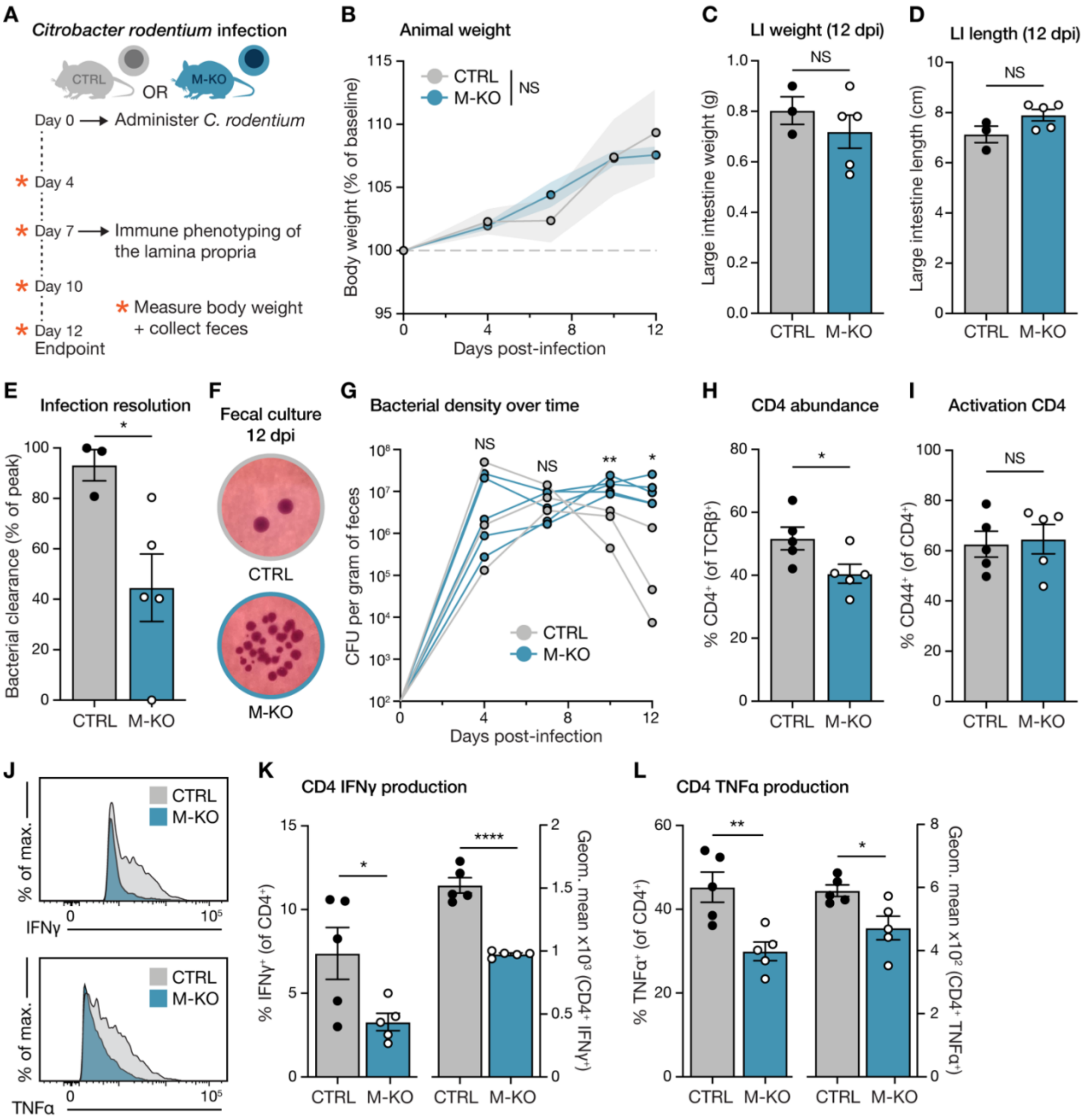


**Figure S5, related to Figure 2: T cell-specific MCAT loss dampens CD4⁺ T cell responses to *Citrobacter rodentium* infection.** (**A**) Experimental setup for infection of CTRL or M-KO mice with *Citrobacter rodentium*. (**B**) Body weights of CTRL and M-KO mice following infection with *C. rodentium.* Data are shown as body weight change over the infection time-course, relative to baseline. Data represent the mean ± SEM ($n$ = 3 CTRL, 5 M-KO). (**C-D**) Quantification of large intestine weight (**C**) and length (**D**) in CTRL and M-KO mice at 12 days post-infection (dpi). Data represent the mean ± SEM ($n$ = 3 CTRL, 5 M-KO). (**E**) Quantification of infection resolution in CTRL and M-KO mice, calculated by comparing fecal bacterial concentrations at 12 dpi to peak levels. Data represent the mean ± SEM ($n$ = 3 CTRL, 5 M-KO). (**F**) Representative images of fecal samples from CTRL and M-KO mice grown on MacConkey agar plates to visualize *C. rodentium* colonies. (**G**) Number of colony forming units (CFU) per gram of feces from CTRL and M-KO mice across the 12 day infection time-course. Individual biological replicates are shown. Dot and line color denotes animal genotype. (**H-I**) Quantification of the percentage of $CD4^+$ (**H**) and $CD4^+$ $CD44^+$ (**I**) T cells in the lamina propria of CTRL and M-KO mice at 7 dpi. Data represent the mean ± SEM (*n* = 5 mice per group). (**J-L**) Representative flow cytometry histograms (**J**) and quantification of IFN-ɣ (**K**) and TNF-α (**L**) production by $CD4^+$ T cells in the lamina propria of CTRL and M-KO mice at 7 dpi. Data are displayed as both the percentage of IFN-$ɣ^+$/TNF-$α^+$ $CD4^+$ cells and the mean fluorescence intensity (MFI) of IFN-ɣ/TNF-α levels in $CD4^+$ cells. Data represent the mean ± SEM (*n* = 5 mice per group). For statistics: **p* > 0.05, ***p* < 0.01, *****p* < 0.0001 by unpaired *t* test with Welch's correction (**C-E, H-I and K-L**), or two-way ANOVA with Tukey's multiple comparisons test (**B and G**), NS denotes not significant.

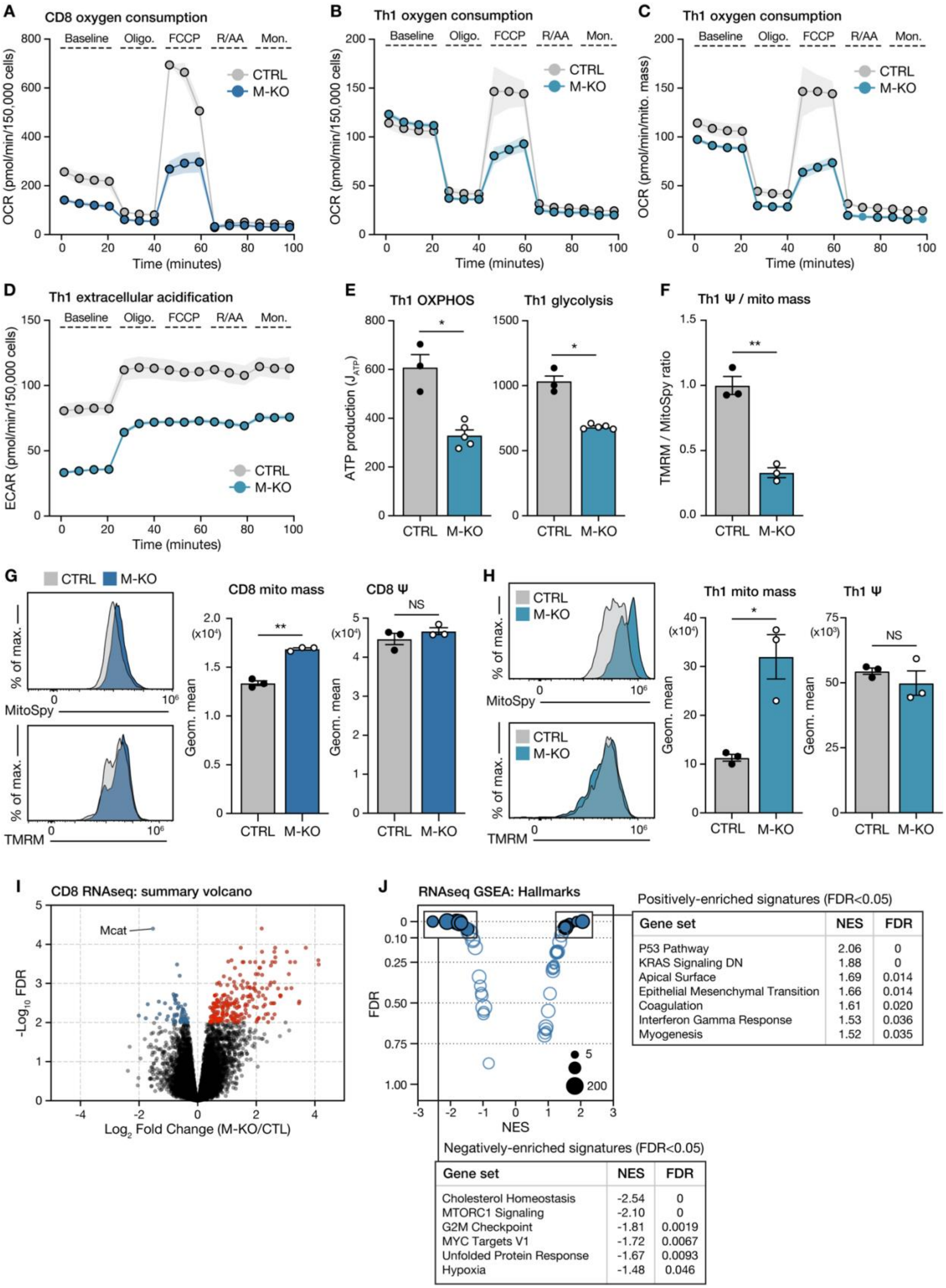


Positively-enriched signatures (FDR<0.05)

| Gene set | NES | FDR |
|---|---|---|
| P53 Pathway | 2.06 | 0 |
| KRAS Signaling DN | 1.88 | 0 |
| Apical Surface | 1.69 | 0.014 |
| Epithelial Mesenchymal Transition | 1.66 | 0.014 |
| Coagulation | 1.61 | 0.020 |
| Interferon Gamma Response | 1.53 | 0.036 |
| Myogenesis | 1.52 | 0.035 |

Negatively-enriched signatures (FDR<0.05)

| Gene set | NES | FDR |
|---|---|---|
| Cholesterol Homeostasis | -2.54 | 0 |
| MTORC1 Signaling | -2.10 | 0 |
| G2M Checkpoint | -1.81 | 0.0019 |
| MYC Targets V1 | -1.72 | 0.0067 |
| Unfolded Protein Response | -1.67 | 0.0093 |
| Hypoxia | -1.48 | 0.046 |

**Figure S6, related to Figure 3: MCAT loss impairs T cell oxygen consumption and bioenergetic capacity.** (**A-D**) Bioenergetic assessment of activated CTRL and M-KO CD8$^+$ T and CD4$^+$ Th1 cells, as quantified using the Seahorse mitochondrial stress test. Shown are the oxygen consumption rate (OCR) of CD8$^+$ T cells normalized to cell number (**A**), the OCR of CD4$^+$ Th1 cells normalized to cell number (**B**), the OCR of CD4$^+$ Th1 cells normalized to mitochondrial content (**C**), the extracellular acidification rate (ECAR) of CD4$^+$ Th1 cells normalized to cell number (**D**). Data show one representative sample per genotype from 3 independent biological replicates, error bars denote SD. (**E**) Calculated maximal ATP production rates derived from oxidative phosphorylation (OXPHOS) and glycolysis in CTRL and M-KO CD4$^+$ Th1 cells. Data represent the mean ± SEM ($n$ = 3 CTRL, 5 M-KO). (**F**) Quantification of mitochondrial membrane potential ($\Psi$) normalized to cellular mitochondrial content in CTRL and M-KO CD4$^+$ Th1 cells. Data represent the mean ± SEM ($n$ = 3 mice per group). (**G and H**) Quantification of mitochondrial mass (MitoSpy) and mitochondrial membrane potential (TMRM) in activated CTRL and M-KO CD8$^+$ T (**G**) and CD4$^+$ Th1 (**H**) cells. Shown are representative histograms (left panels) and mean fluorescence intensity values (right panels) of each stain individually. Data represent the mean ± SEM ($n$ = 3 mice per group). (**I**) Volcano plot summarizing differentially-expressed genes in activated CTRL and M-KO CD8$^+$ T cells, according to RNAseq data. Data are shown are relative to CTRL (Log2 fold change, M-KO/CTRL), colored dots denote FDR < 0.01 ($n$ = 6 mice per group). (**J**) RNAseq gene set enrichment analysis (GSEA, MSigDB Hallmarks) showing GO terms that enriched in M-KO versus CTRL CD8$^+$ Th1 cells. Bubble plot displays the normalized enrichment score (NES) and false discovery rate (FDR) for all analyzed signatures; bubble size equates to gene list size. All significantly regulated signatures (FDR < 0.05) are highlighted and listed alongside their NES and FDR values. For statistics: *$p$ > 0.05, **$p$ < 0.01 by unpaired $t$ test with Welch's correction (**E-H**), NS denotes not significant.

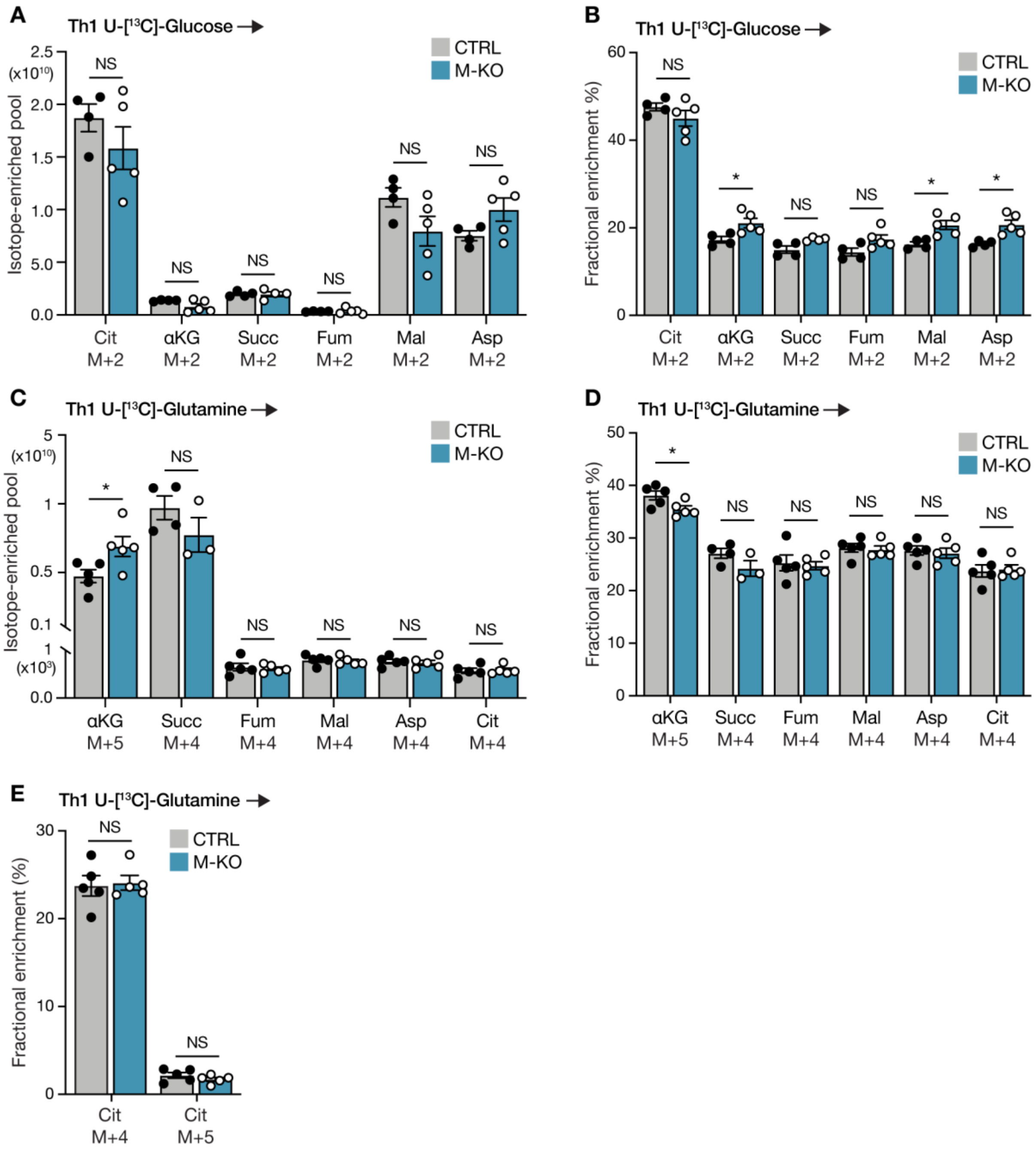


**Figure S7, related to Figure 3: Glucose and glutamine utilization in CD4$^+$ Th1 cells is largely unchanged by MCAT loss.** (**A-E**) Contribution of [U-$^{13}$C]-glucose (**A and B**) and -glutamine (**C-E**) to TCA cycle intermediates and aspartate in activated CTRL and M-KO CD4$^+$ Th1 cells following a 4h trace. Data shown represent the total abundance of the indicated mass isotopologues derived from the labeled tracer (**A and C**), or fractional enrichment of the labeled tracer in the indicated mass isotopologues (**B, D and E**). Error bars denote SEM (n = 5 mice per group per tracer). For statistics: *$p$ > 0.05 by unpaired $t$ test with Welch's correction (**A-D**), or two-way ANOVA with Tukey's multiple comparisons test (**E**), NS denotes not significant.

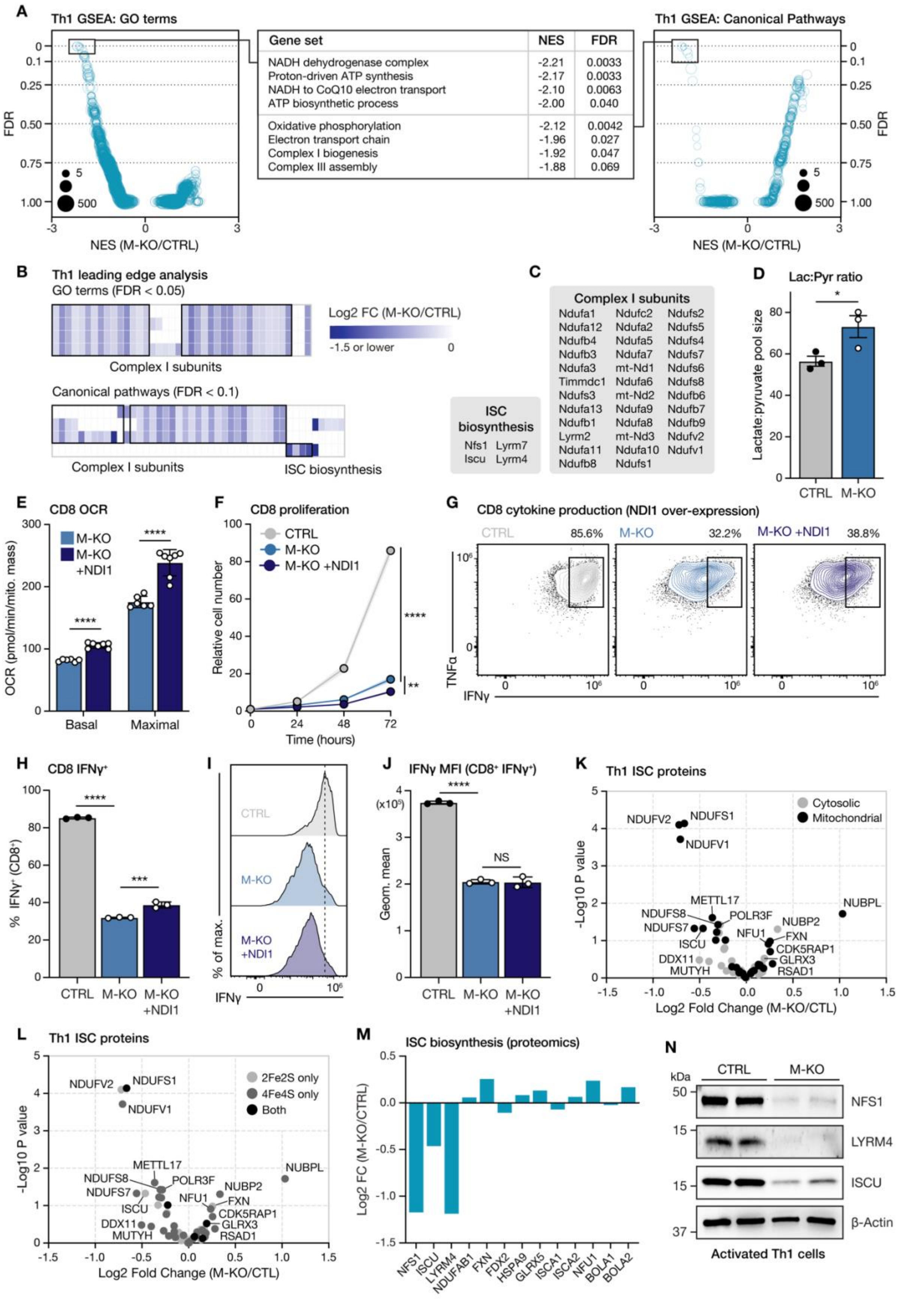
A
Th1 GSEA: GO terms
FDR
NES (M-KO/CTRL)
Gene set | NES | FDR
NADH dehydrogenase complex | -2.21 | 0.0033
Proton-driven ATP synthesis | -2.17 | 0.0033
NADH to CoQ10 electron transport | -2.10 | 0.0063
ATP biosynthetic process | -2.00 | 0.040
Oxidative phosphorylation | -2.12 | 0.0042
Electron transport chain | -1.96 | 0.027
Complex I biogenesis | -1.92 | 0.047
Complex III assembly | -1.88 | 0.069
Th1 GSEA: Canonical Pathways
B
Th1 leading edge analysis
GO terms (FDR < 0.05)
Log2 FC (M-KO/CTRL)
-1.5 or lower
Complex I subunits
Canonical pathways (FDR < 0.1)
Complex I subunits
ISC biosynthesis
C
ISC biosynthesis
Nfs1 Lyrm7
Iscu Lyrm4
Complex I subunits
Ndufa1 Ndufc2 Ndufs2
Ndufa12 Ndufa2 Ndufs5
Ndufb4 Ndufa5 Ndufs4
Ndufb3 Ndufa7 Ndufs7
Ndufa3 mt-Nd1 Ndufs6
Timmdc1 Ndufa6 Ndufs8
Ndufs3 mt-Nd2 Ndufb6
Ndufa13 Ndufa9 Ndufb7
Ndufb1 Ndufa8 Ndufb9
Lyrm2 mt-Nd3 Ndufv2
Ndufa11 Ndufa10 Ndufv1
Ndufb8 Ndufs1
D
Lac:Pyr ratio
Lactate:pyruvate pool size
CTRL
M-KO
E
CD8 OCR
M-KO
M-KO +NDI1
OCR (pmol/min/mito. mass)
Basal
Maximal
F
CD8 proliferation
CTRL
M-KO
M-KO +NDI1
Relative cell number
Time (hours)
G
CD8 cytokine production (NDI1 over-expression)
CTRL 85.6%
M-KO 32.2%
M-KO +NDI1 38.8%
TNFα
IFNγ
H
CD8 IFNγ+
% IFNγ+ (CD8+)
CTRL
M-KO
M-KO +NDI1
I
CTRL
M-KO
M-KO +NDI1
% of max.
IFNγ
J
IFNγ MFI (CD8+ IFNγ+)
(x10⁵)
Geom. mean
NS
CTRL
M-KO
M-KO +NDI1
K
Th1 ISC proteins
Cytosolic
Mitochondrial
-Log10 P value
Log2 Fold Change (M-KO/CTL)
NDUFV2
NDUFS1
NDUFV1
METTL17
NDUFS8
POLR3F
NDUFS7
NUBP2
NFU1
FXN
ISCU
CDK5RAP1
DDX11
GLRX3
MUTYH
RSAD1
NUBPL
L
Th1 ISC proteins
2Fe2S only
4Fe4S only
Both
-Log10 P value
Log2 Fold Change (M-KO/CTL)
M
ISC biosynthesis (proteomics)
Log2 FC (M-KO/CTRL)
NFS1
ISCU
LYRM4
NDUFAB1
FXN
FDX2
HSPA9
GLRX5
ISCA1
ISCA2
NFU1
BOLA1
BOLA2
N
kDa
CTRL
M-KO
NFS1
LYRM4
ISCU
β-Actin
Activated Th1 cells

**Figure S8, related to Figure 4: Disrupted complex I formation and iron-sulfur cluster biosynthesis are central features of MCAT-knockout T cells.** (**A**) Gene set enrichment analysis (GSEA, MSigDB C2 and C5) showing GO terms and canonical pathways that are enriched in M-KO versus CTRL $CD4^+$ Th1 T cells. Bubble plots display the normalized enrichment score (NES) and false discovery rate (FDR) for all analyzed signatures; bubble size equates to gene list size. All significantly regulated signatures are listed (middle table) alongside their NES and FDR values. (**B**) Heatmap of protein expression (log2 fold change, M-KO/CTRL) for leading-edge genes in all signatures with significant negative enrichment in M-KO $CD4^+$ Th1 versus CTRL T cells. (**C**) List of electron transport chain (ETC) subunits and iron-sulfur cluster (ISC) biosynthesis proteins identified by leading edge analysis in **B**. ISC-containing ETC complex I subunits are underlined. (**D**) Ratio of total lactate and pyruvate pool sizes in activated CTRL and M-KO $CD8^+$ T cells as measured by metabolomics. Data represent the mean ± SEM (*n* = 3 mice per group). (**E-J**) M-KO mice were crossed with *Ndi1*-expressing knock-in mice to generate M-KO and 'M-KO +NDI1' $CD8^+$ T cells, which express an alternative NADH dehydrogenase found in *Saccharomyces cerevisiae*. (**E**) Basal and maximal oxygen consumption of M-KO and M-KO +NDI1 $CD8^+$ T cells, as measured using the Seahorse mitochondrial stress test. Data represent the mean ± SD (*n* = 6 technical replicates using cells pooled from 2 mice per genotype). (**F**) Proliferation of CTRL, M-KO and M-KO +NDI1 $CD8^+$ T cells cultured in IL-2 (50 U/mL). Naïve cells activated for 3d in vitro using plate-bound anti-CD3 and anti-CD28 antibodies; the number of viable T cells are displayed. Data represent the mean ± SD (*n* = 3 technical replicates using cells pooled from 2 mice per genotype). (**G-J**) Cytokine production of activated CTRL, M-KO and M-KO +NDI1 $CD8^+$ T cells following a 4h PMA/ionomycin stimulation. Shown are representative flow plots of IFN-γ versus TNF-α expression (**G**) the percentage IFN-$\gamma^+$ $CD8^+$ cells (**H**), representative histograms of IFN-γ expression in $CD8^+$ T cells (**I**), and the MFI of IFN-γ levels in IFN-$\gamma^+$ $CD8^+$ T cells (**J**). Data represent the mean ± SD (*n* = 3 technical replicates using cells pooled from 2 mice per genotype). (**K and L**) Volcano plots displaying the protein levels of known ISC proteins in CTRL and M-KO $CD4^+$ Th1 cells. Protein abundance is shown relative to CTRL (log2 fold change, M-KO/CTRL); dot color denotes cytosolic versus mitochondrial localization (**K**) or type of ISC involved (**L**). (**M**) Waterfall plot displaying the protein level of ISC biosynthesis machinery in CTRL and M-KO $CD4^+$ Th1 cells. Data are expressed as relative to CTRL (combined from *n* = 4 CTRL, 3 M-KO). (**N**) Immunoblot of NFS1, LYRM4, ISCU and β-Actin protein levels in activated CTRL and M-KO $CD8^+$ T cells.

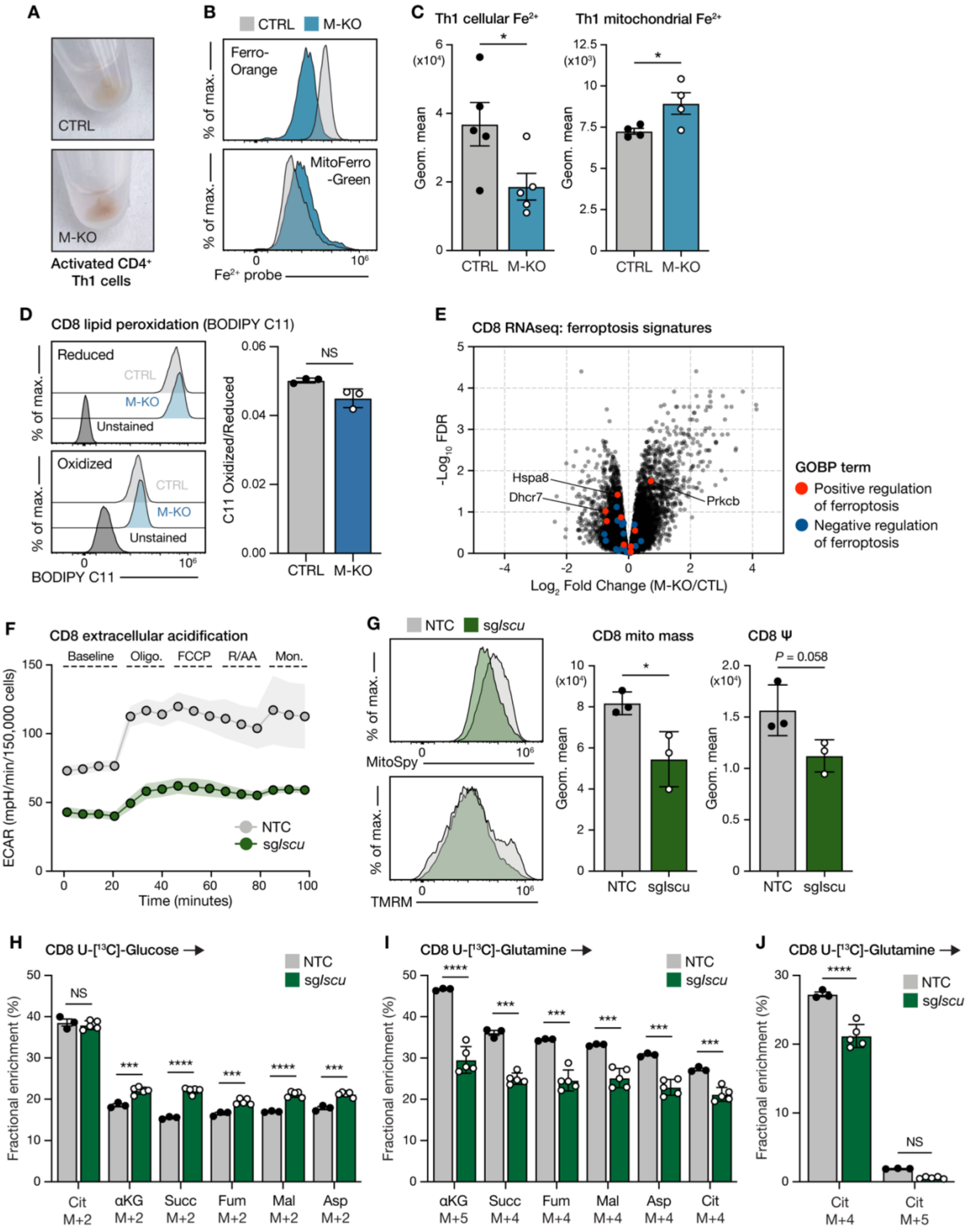


**Figure S9, related to Figure 5: mtFAS supports effector T cell function via regulation of ISC synthesis.** (**A**) Representative pictures of the pellets from activated CTRL and M-KO CD4+ Th1 cell cultures. (**B-C**) Representative flow cytometry histograms (**B**) and quantification of cellular (Ferro-Orange) and mitochondrial (MitoFerro-Green) free iron ($Fe^{2+}$) levels in CTRL and M-KO $CD4^+$ Th1 cells (**C**). Cells were activated in vitro using plate-bound anti-CD3 and anti-CD28 antibodies for 3d; data represent the mean ± SEM (*n* = 5 mice per group). (**D**) Representative histograms (left) and quantified ratio (right) of oxidized and reduced BODIPY C11 fluorescence; an indicator of lipid peroxidation Data represents the mean ± SEM following a 30m incubation (*n* = 3 per group). (**E**) Volcano plot summarizing differentially-expressed genes in activated CTRL and M-KO $CD8^+$ T cells, according to RNAseq data. Data are shown are relative to CTRL (Log2 fold change, M-KO/CTRL), colored dots denote genes associated with either positive (red dots) or negative (blue dots) regulation of ferroptosis according to MSigDB GOBP signatures (*n* = 6 mice per group). (**F**) Extracellular acidification rate (ECAR) of activated wild-type (NTC) and *Iscu*-deleted (sg*Iscu*) $CD8^+$ T cells, as quantified using the Seahorse mitochondrial stress test. Data show one representative technical replicate (*n* = 3 technical replicates per group using cells pooled from 3 mice), error bars denote SD. (**G**) Quantification of mitochondrial mass (MitoSpy) and mitochondrial membrane potential (TMRM) in activated wild-type (NTC) and *Iscu*-deleted $CD8^+$ T cells. Shown are representative histograms (left) and mean fluorescence intensity values (right) of each stain individually. Data represent the mean ± SD (*n* = 3 technical replicates per group using cells pooled from 3 mice). (**H-J**) Contribution of [U-$^{13}$C]-glucose (**H**) and -glutamine (**I and J**) to TCA cycle intermediates and aspartate in wild-type (NTC) and *Iscu*-deleted $CD8^+$ T cells. Data shown represent the fractional enrichment of the labeled tracer in the indicated mass isotopologues. Error bars denote SD (*n* = 3 NTC and 5 sg*Iscu* technical replicates using cells pooled from 3 mice). For statistics: *$p > 0.05$, ***$p < 0.001$, ****$p < 0.0001$ by unpaired *t* test with Welch's correction (**C, D and G**), one-way ANOVA with Tukey's multiple comparisons test (**H and I**), or two-way ANOVA with Tukey's multiple comparisons test (**J**), NS denotes not significant.

Supplemental Figures (Preston et al.)

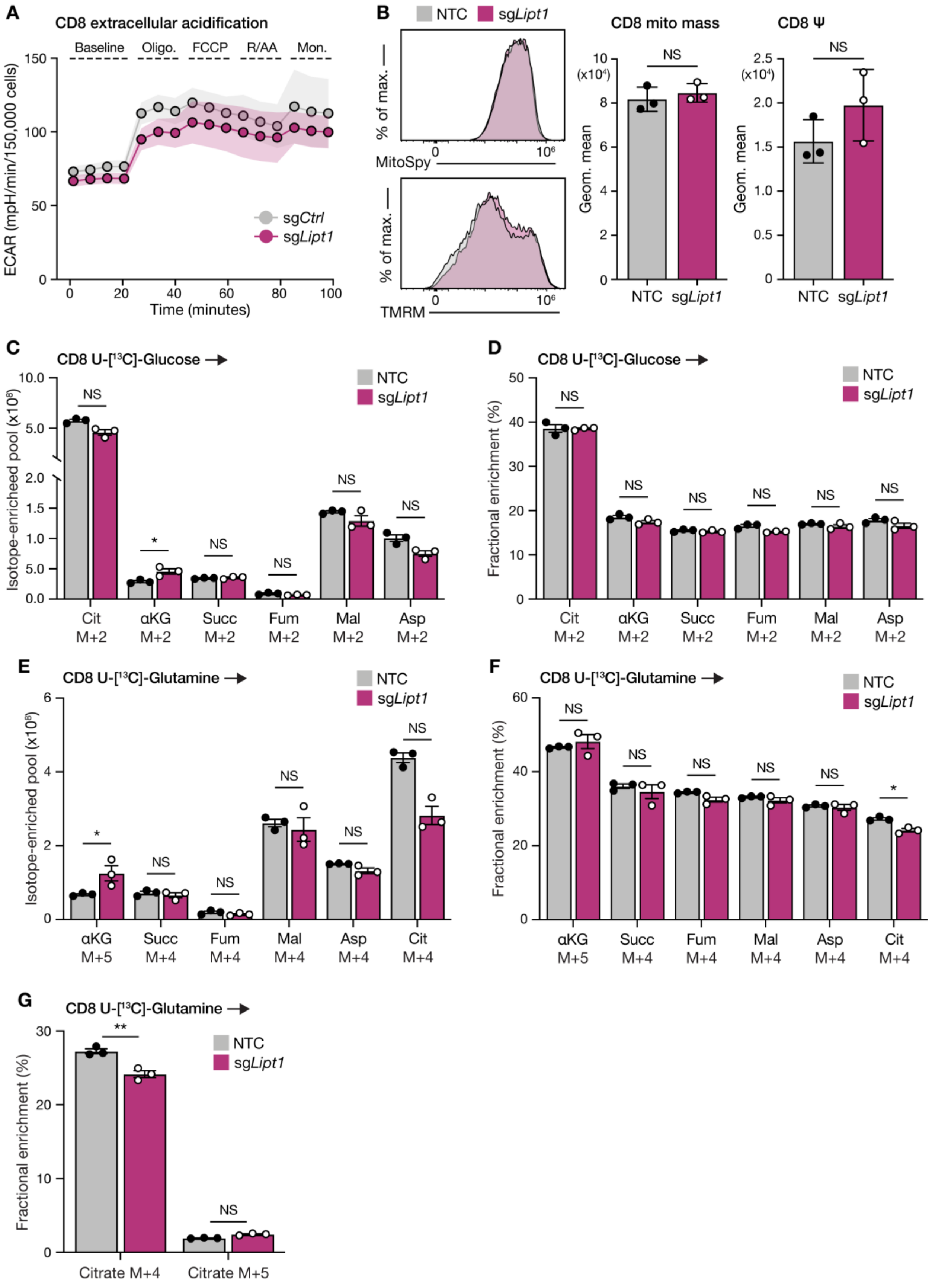

**Figure S10, related to Figure 6: Bioenergetic and metabolic characterization of *Lipt1*-deleted CD8$^{+}$ T cells.** (**A**) Extracellular acidification rate (ECAR) of activated wild-type (NTC) and *Lipt1*-deleted (sg*Lipt1*) CD8$^{+}$ T cells, as quantified using the Seahorse mitochondrial stress test. Data show one representative technical replicate ($n$ = 3 technical replicates per group using cells pooled from 3 mice), error bars denote SD. (**B**) Quantification of mitochondrial mass (MitoSpy) and mitochondrial membrane potential (TMRM) in activated wild-type (NTC) and *Lipt1*-deleted CD8$^{+}$ T cells. Shown are representative histograms (left) and mean fluorescence intensity values (right) of each stain individually. Data represent the mean ± SD ($n$ = 3 mice per group). (**C-G**) Contribution of [U-$^{13}$C]-glucose (**C and D**) and -glutamine (**E-G**) to TCA cycle intermediates and aspartate in activated wild-type (NTC) and *Lipt1*-deleted CD8$^{+}$ T cells following a 4h trace. Data shown represent the total abundance of the indicated mass isotopologues derived from the labeled tracer (**C and E**), or fractional enrichment of the labeled tracer in the indicated mass isotopologues (**D, F and G**). Error bars denote SD ($n$ = 3 technical replicates per tracer using cells pooled from 3 mice). For statistics: *$p > 0.05$, **$p > 0.01$ by unpaired $t$ test with Welch's correction (**B**), one-way ANOVA with Tukey's multiple comparisons test (**C-F**), or two-way ANOVA with Tukey's multiple comparisons test (**G**), NS denotes not significant.

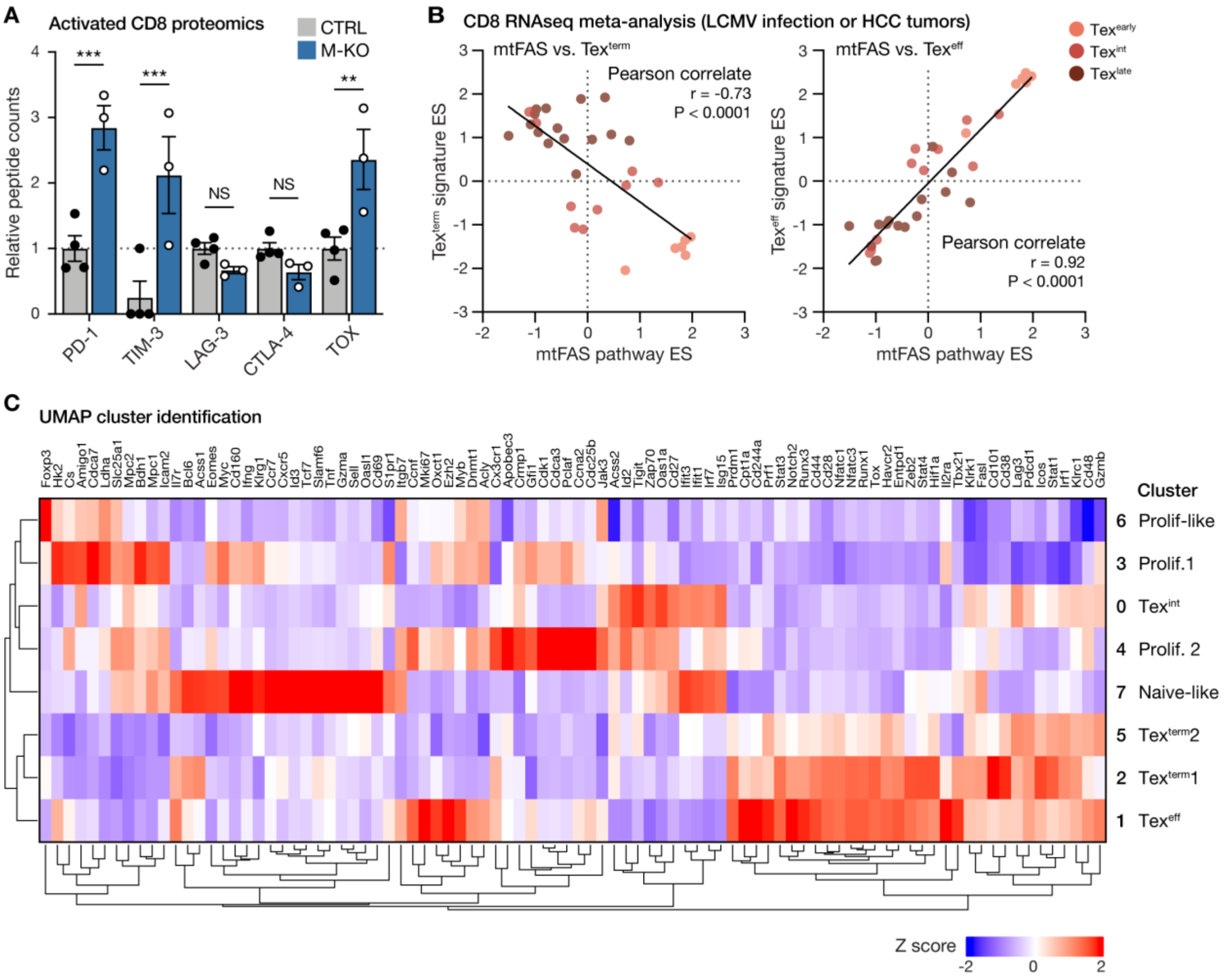


**Figure S11, related to Figure 7: mtFAS deficiency promotes CD8⁺ T cell exhaustion.** (**A**) Relative peptide counts of indicated inhibitory receptors (PD-1, TIM-3, LAG-3, and CTLA-4) and TOX) in CTRL and M-KO CD8⁺ T cells 3d following anti-CD3 and anti-CD28 activation, as detected by proteomics. Data are shown as relative to levels in CTRL cells (dotted line) and represent the mean ± SEM (*n* = 4 for CTRL, 3 for M-KO). (**B**) Single-sample GSEA of RNA-seq data from early-, intermediate-, and late-exhausted CD8⁺ T cells (denoted by dot color) isolated from mice infected with LCMV or bearing liver tumors. Pearson rank-order correlations of enrichment scores are shown for the mtFAS pathway against a signature of terminally exhausted ($Tex^{term}$, left panel) or effector-like exhausted ($Tex^{eff}$, right panel) CD8⁺ T cells. Solid black lines show linear regression. *P* and *r* values are indicated. (**C**) Heatmap summarizing the transcript expression data used to annotate population clusters on the UMAP in **Fig. 7A**. Data are shown as row-normalized Z scores.
